# Allele-resolved single-cell multi-omics uncovers the dynamics and transcriptional kinetics of X-chromosome upregulation

**DOI:** 10.1101/2021.07.14.452323

**Authors:** Antonio Lentini, Huaitao Cheng, JC Noble, Natali Papanicolaou, Christos Coucoravas, Nathanael Andrews, Qiaolin Deng, Martin Enge, Björn Reinius

## Abstract

X-chromosome inactivation (XCI) and upregulation (XCU) are the major opposing chromosome-wide modes of gene regulation that collectively achieve dosage compensation in mammals, but the regulatory link between the two remains elusive. Here, we use allele-resolved single-cell RNA-seq combined with chromatin accessibility profiling to finely dissect the separate effects of XCI and XCU on RNA levels during mouse development. We uncover that balanced X dosage is flexibly attained through expression tuning by XCU in a sex- and lineage-specific manner along varying degrees of XCI and across developmental and cellular states. Male blastomeres achieve XCU upon zygotic genome activation while females experience two distinct waves of XCU, upon imprinted- and random XCI, and ablation of *Xist* impedes female XCU. Contrary to widely established models of mammalian dosage compensation, naïve female embryonic cells carrying two active X chromosomes do not exhibit upregulation but express both alleles at basal level, yet collectively exceeding the RNA output of a single hyperactive allele. We show, *in vivo* and *in vitro*, that XCU is kinetically driven by X-specific modulation of transcriptional burst frequency, coinciding with increased compartmentalization of the hyperactive allele. Altogether, our data provide unprecedented insights into the dynamics of mammalian XCU, prompting a revised model of the chain in events of allelic regulation by XCU and XCI in unitedly achieving stable cellular levels of X-chromosome transcripts.

## Introduction

In therian mammals, the X chromosome is present as two copies in females but only one in males. Correct balance of gene dosage is vital for homeostasis and normal cell function, and two major X-chromosome-wide mechanisms ensure balanced genomic expression^1^. XCU resolves X-to-autosomal gene-dose imbalances in males by hyperactivation of the X chromosome whereas XCI silences one X allele in females, equalizing expression between the sexes^1^. It was first proposed in the 1960s that two-fold expression upregulation through XCU evolved as a first step to compensate for degradation and gene loss of the Y chromosome where evolution of XCI followed as a second step to avoid the lethal dose of two hyperactive X alleles in females^2^. This still stands as the prevailing evolutionary hypothesis^3,4^. However, despite being central for the regulatory model of dosage compensation, the developmental sequence of upregulation and inactivation has remained elusive. The question of XCU timing and dynamics is further perplexed as key model species such as rodents experience two waves of XCI^5,6^ which would in theory expose female embryos to a lethal X dosage twice. Indeed, while the timeline and molecular mechanisms of XCI are well characterized and known to be linked to the pluripotency state^7^, the dynamics and regulation of XCU are largely undetermined. Previous studies have approximated XCU primarily by relative measurements between non-allelic total expression levels of X and autosomes in steady-state^8–13^ which is convoluted in the presence of confounding allelic processes, such as XCI. It is therefore unsurprising that reports on establishment, maintenance, and potential reversal of XCU have been conflicting^8,14,15^. Disentangling the isolated effects of XCU and XCI is not trivial and would require quantitative gene expression measurements at cellular and allelic resolution, only recently enabled by allele-resolved single-cell RNA-seq (scRNA-seq). Additionally, this requires careful computational dissection of stochastic allelic processes affecting expression measurements at the cellular level^16^, including the effects of stochastic transcriptional bursting on allelic expression as well as random XCI (rXCI), progressing heterogeneously in individual cells.

Here, we uncover the XCU dynamics in mouse at cellular and allelic resolution throughout embryonic stem cell priming *in vitro* as well as early embryonic development *in vivo*. Surprisingly, our data reveal that XCU is neither a constitutive state of hyper transcription of the X chromosome, nor established as a discrete regulatory event, but is a flexible process that quantitatively tunes RNA synthesis proportionally to the output of the second X allele via modulation of transcriptional bursting. We demonstrate sex- and lineage-specific initiation and maintenance of XCU in conjunction with gene-regulatory events that would otherwise leave expression dosage unbalanced. By combining allele-resolved scRNA-seq and scATAC-seq (single-cell Assay for Transposase-Accessible Chromatin using sequencing) in the same cells during XCU establishment, we characterized the epigenetics state of XCU. Finally, building on these new insights, we provide a revised model of dosage compensation in the early mammalian embryogenesis, bridging the allele-specific dynamics of XCU and XCI.

## Results

### Female naïve mESCs do not exhibit X-chromosome upregulation

Exit from pluripotency in mouse embryonic stem cells (mESCs) is accompanied by rXCI in females^17^. We modelled this process by differentiating naïve mESCs of mixed genetic background (C57BL6/J × CAST/EiJ) cultured under 2i [Gsk3+MEK inhibition] condition towards primed epiblast stem cells (EpiSCs; Activin A + FGF2) for up to 7 days *in vitro*, during which single cells were captured (**Fig 1a, Methods**). These F1 hybrid cells carry a high density of parental-origin-specific single-nucleotide polymorphisms which we leveraged using sensitive full-length scRNA-seq (Smart-seq3^18^) to obtain allelic information across transcripts and RNA counts by unique molecular identifiers (UMIs) (**Extended Data Fig. 1a**). After quality filtering, we captured allelic expression of up to 576 X-linked and 18,043 autosomal genes in 687 cells, highlighting that our data provided near genome-wide allelic resolution and the sensitivity to study allelic regulation on the gene-level in single cells.

**Figure 1.**
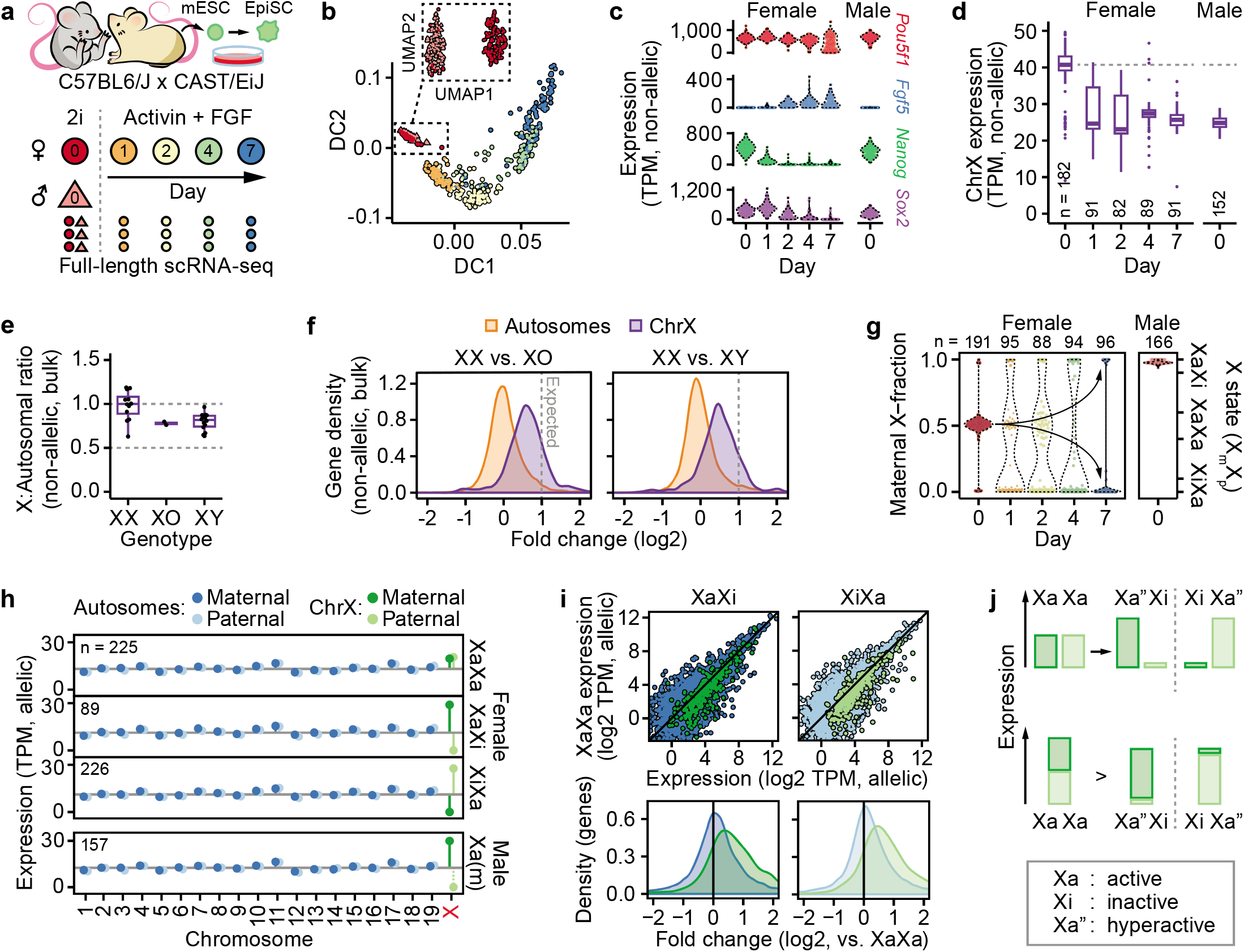
The state of two active X chromosomes lack X-upregulation. **a.** Schematic overview of experimental setup. C57BL6/J × CAST/EiJ F1 hybrid male and female mESCs were maintained under 2i conditions and female mESCs were primed to EpiSCs using Activin A and FGF2 for up to 7 days to induce X-inactivation (XCI). Cells were collected at conditions and days of priming as indicated and subjected to scRNA-seq using Smart-seq3. **b.** Diffusion map of mESC priming towards EpiSCs using the top 1,000 variable genes for 687 cells. Inset shows UMAP dimensions for male (triangle) and female (circle) mESC in 2i conditions. Cell colors as denoted in panel a. **c.** Violin plots of marker gene expression of pluripotency and differentiation markers *Pou5f1, Fgf5, Nanog* and *Sox2* along EpiSC priming. **d.** Boxplots showing total expression of chrX per cell, sex and timepoint along EpiSC priming (n = 1,158–1,337 genes). **e.** X:Autosomal ratios for bulk RNA-seq of mESCs of XX, XO (Turner syndrome) and XY genotypes (n = 409–522 chrX genes). N bulk RNA-seq libraries: XO (2), XY (14), XX (12). N genes: autosomal (10,961), X (405). **f.** Density plot of gene-wise expression fold changes in bulk RNA-seq from female mESCs carrying two X-chromosomes (XX) relative to female mESCs lacking one X-chromosome copy (XO) or male mESCs (XY). Dashed lines indicate expected two-fold expression fold change, n = same as in e. **g.** Violin plots of fraction maternal X-chromosome expression in cells (dots) along EpiSC priming. Female cells were classed according to allelic X-chromosome expression bias, i.e. XCI state (right y-axis). **h.** Allele-resolved expression per cell for all chromosomes grouped by sex and XCI status, for autosomes (blue; n = 15,683–16,543 genes) and X (green; n = 508–518), plotted as median ± 95% confidence interval. **i.** Expression scatter plots of the same active allele for XaXa and X-inactive states (XaXi or XiXa; C57 and CAST allele active, respectively) in female cells (left) and density plots of allelic expression relative to the XaXa state (right). Blue: Autosomal genes (n = 13,696–14,845), green: X-linked genes (n = 383–433). **j.** Schematic summarization of key finding and notation of transcriptional states. Female naïve mESCs with of two active X chromosomes (XaXa) lack XCU. As XCI inactivates one allele (Xi) the remaining becomes transcriptionally hyperactive (Xa”). Biallelic RNA output of XaXa exceeds that of the monoallelic Xa” state.

The mESC-to-EpiSC transition triggered distinct expression changes accompanied by the loss of pluripotency factors (*e.g. Sox2* & *Nanog*) and induction of lineage-specific factors (*e.g. Fgf5* & *Krt18*) together with related pathways (**Fig. 1b-c, Extended Data Fig. 1b-c and Supplementary Table 1**), signifying successful stem cell priming. As expected, female naïve mESCs, carrying two active X alleles (XaXa state), demonstrated an elevated total X-gene expression dosage that diminished upon exit from pluripotency (**Fig. 1d**). This has previously been attributed to the silencing of one out of two hyperactive X alleles through XCI^10^. We confirmed the elevated X dosage in naïve female XX mESCs compared to both male XY and female XO (Turner syndrome) mESCs in bulk RNA-seq^19,20^ (**Fig. 1e**). At the same time, we noticed that the female XX mESC expression was less than the two-fold higher expected relative to XY and XO cells (median XY = 1.42 and XO = 1.54 fold, relative to XX) if comparing a biallelic hyperactive XaXa state to a single hyperactive-X allele in XY/XO cells (**Fig. 1f**). To dissect X dosage to the allelic resolution, we stratified our single-cell data according to XCI status inferred from X-linked allelic expression [fraction of expression maternal/(maternal + paternal)], into biallelic and monoallelic X-chromosome states (XaXa, and XaXi / XiXa, respectively) (**Fig. 1g and Extended Data Fig. 1d**) indeed confirming decreased total X-linked expression upon XCI in female cells (**Extended Data Fig. 1e**).

Surprisingly, when resolved onto the separate alleles, X-linked expression in XaXa cells was on par with autosomal levels for each allele whereas female X-inactive states (XaXi/XiXa) and male (XY) cells exhibited distinct upregulation of the single active X-chromosome copy (**Fig. 1h**). Intriguingly, the lack of XCU in XaXa state implies that XCI does not silence a hyperactive X allele, which is conceptually distinct from what is widely believed and modelled^3,4^. Notably, upregulation was observed X-chromosome-wide (**Extended Data Fig. 1f**) and female XaXa cells consistently lacked XCU regardless of days of EpiSC priming (**Extended Data Fig. 1g**), ruling out sporadic effects related to the differentiation process. To independently validate these findings, we reanalyzed recently published 3’ UMI-tagged allele-specific scRNA-seq data of 2i withdrawal in mESCs^21^. Under these conditions, XCI is spontaneously initiated at a considerably lower rate compared to EpiSC priming^17^, allowing us to observe the effect of biallelic versus monoallelic X expression over a wider timespan (**Extended Data Fig. 1h**). In agreement with our EpiSC priming results, we observed that total X-linked expression aggregated across both alleles was consistently higher in XaXa cells than those undergoing XCI at all timepoints, yet lack of biallelic XCU in the XaXa state (**Extended Data Fig. 1i-j**).

Our finding that two moderately expressed X alleles (XaXa state) achieve a higher total expression dosage than a single hyperactive X allele suggests that XCU does not attain complete compensation at the transcript-count level (*i.e*. less than two-fold upregulation). Indeed, we calculated relative gene-wise expression of the same active allele transitioning from XaXa into XaXi state (**Fig. 1i**) which demonstrated a pronounced shift in X-linked expression yet with fold-changes below two (median X_C57_ = 1.62, P_C57_ = 1.41 × 10^−40^ and median X_CAST_ = 1.67, P_CAST_ = 5.72 × 10^−48^, Wilcoxon signed-rank test). While these findings conceptually conflict the assumption that XCI acts on a biallelically hyperactive XaXa state^3,4^, the scenario remains compatible with the notion that expression dosage of two active X alleles blocks or delays differentiation of female mESCs^22^ as the total X dosage is reduced when female mESCs transition from naïve to primed pluripotency (XaXa → hyperactive Xa’’Xi state, where’’ denotes increased transcriptional activity) (**Fig. 1j**).

In summary, our results reveal the important finding that the two active X alleles in female naïve mESCs lack X-upregulation, whereas hyperactivation of transcription (<2-fold) is present in cellular states with a single active X-chromosome copy.

### Two distinct waves of X-chromosome upregulation during early mouse development

Naïve mESCs are derived from, and mimic, the inner cell mass (ICM) of pre-implantation embryos. XCU was previously proposed to be present prior to ICM formation^8,14,15^ but the lack of hyperactivation in naïve female mESCs we observed here suggested that XCU might be turned on- and off during embryonic development. To investigate this further, we leveraged the large full-length allele-level scRNA-seq datasets we recently generated across early murine embryogenesis (**Fig. 2a-b**)^6,17,23^. These data cover key developmental timepoints from the oocyte/zygote up until gastrulation and span the major known embryonic events of X-chromosome regulation^1^, *i.e*. imprinted XCI (iXCI), X-chromosome reactivation (XCR), and rXCI (**Fig. 2c**).

**Figure 2.**
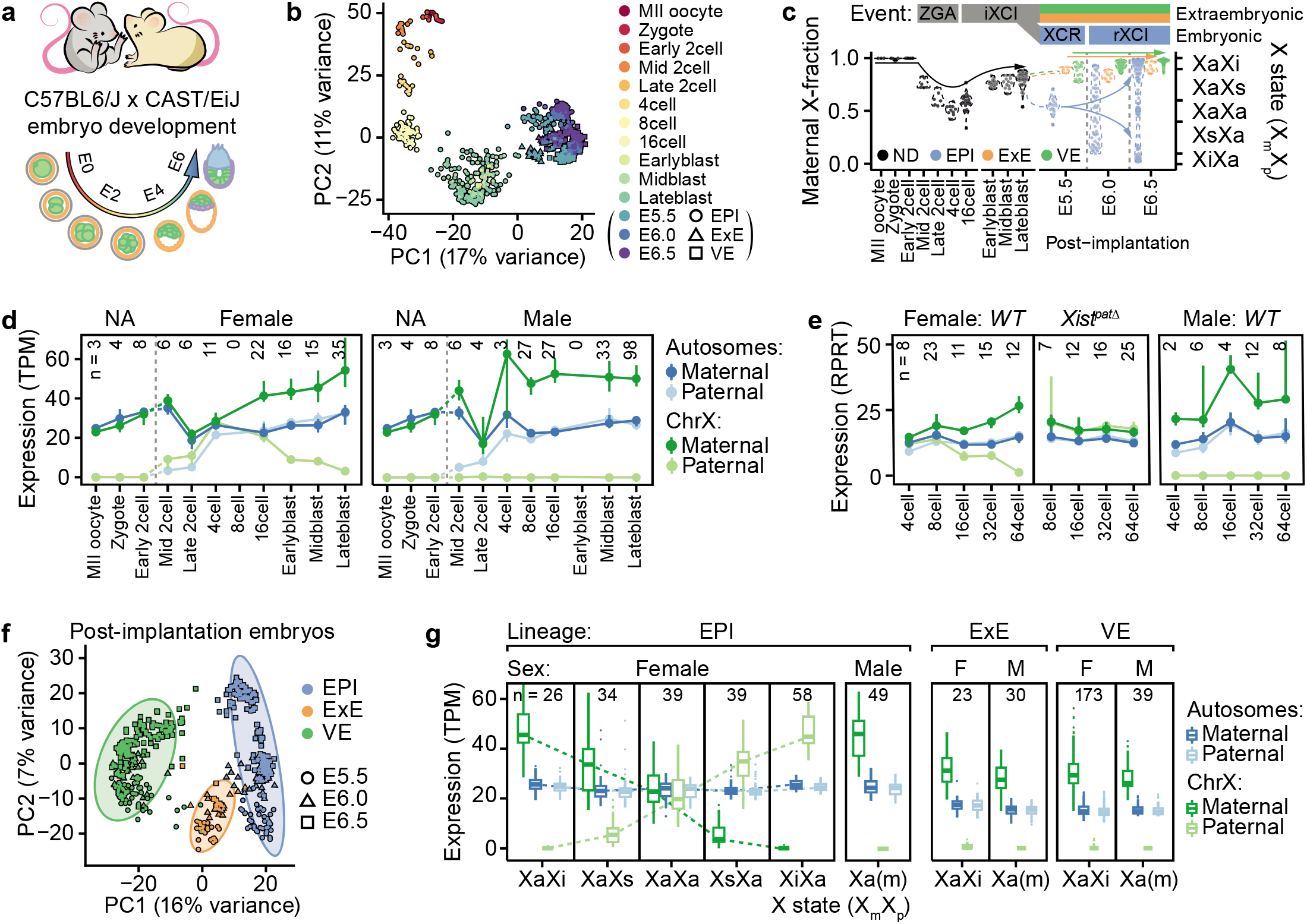
X-upregulation dynamics during early embryonic development in vivo. **a.** Schematic overview of allele-specific scRNA-seq datasets from C57BL6/J × CAST/EiJ F1 hybrid embryos ranging from Zygote to post-implantation stages. **b.** PCA plot for the top 2,000 HVGs of 834 cells across early murine embryonic development. **c.** Violin plots of maternal fraction X-chromosome expression in cells (dots) across the first week of female mouse embryo development (x-axis), with known key events of allelic and X-chromosome regulation indicated above; ZGA (mid-2- to 4-cell stage), iXCI (initiating at 8- to 16-cell stage), XCR (epiblast-specific, late blastocyst to early implantation) and rXCI (epiblast-specific following XCR). Past ZGA, the XCI states of female cells were inferred according to allelic X-chromosome expression bias (right y-axis). Cell-lineage assignment: not determine (ND), epiblast (EPI), extraembryonic ectoderm (ExE), visceral endoderm (VE). **d.** Allele-resolved expression during pre-implantation development for autosomes (blue; n = 11,164–13,347 genes) and chrX (green; n = 323–437 genes) in cells of female and male embryos. Data shown as median ± 95% confidence interval. **e.** Allele-resolved expression for wild-type (WT) or female Xist^patΔ^ knockout embryo cells along initiation of iXCI for autosomes (blue; 1,712–7,908 genes) and chrX (green; 36–248). Legend as in panel d., RPRT = Reads Per Retro-Transcribed length per million mapped reads. **f.** PCA plot for the top 2,000 HVGs of 510 post-implantation embryo cells grouped by lineage. **g.** Boxplots of allelic gene expression levels in post-implantation cells grouped by sex and lineage. Female epiblast cells are further grouped by rXCI state. Shown for autosomes (blue; n = 14,941–17,643 genes) and chrX (green; n = 597–713).

Allelic expression was balanced for X and autosomes in mature (MII) oocytes up until 2-cell stages where mRNAs originate only from the maternal genome. However, around completion of zygotic genome activation (ZGA) where biallelic autosomal transcription is achieved (~4-cell stage) XCU was specifically observed in male cells (**Fig. 2d**). Conversely, female 4-cell embryos exhibited biallelic XaXa expression that lacked XCU recapitulating our results in naïve mESCs, and maternal-specific XCU was first detected around the 8-16-cell stage (**Fig. 2d**), notably coinciding with iXCI on the paternal allele.

Importantly, these sex-specific temporal dynamics closely followed events that would otherwise result in imbalanced chromosomal dosage, suggesting that XCU primarily acts in response to other X-chromosome-dosage imbalances. To test this hypothesis, we analyzed XCU in allele-resolved scRNA-seq data from *Xist^patΔ^* knockout embryos genetically designed to lack iXCI^14^. Indeed, female 16-to-64-cell knockout embryos did not initiate XCU (**Fig. 2e**), indicating that its transcriptional hyperactivity is initiated as a response to imbalanced dosage and not the inverse. XCU was further maintained in both sexes, along with iXCI in females, as wild-type embryos developed into pre-implantation blastocysts (**Fig. 2d**). Because lineage specification commences during blastocyst development, we classified cells into epiblast (EPI), trophectoderm (TE) and primitive endoderm (PrE) lineages (**Extended Data Fig. 2a-b, Methods**) and investigated XCU in each trajectory, which revealed XCU to be present in all late-blastocyst lineages, including EPI cells prior to XCR (**Extended Data Fig. 2c**), again showing that XCU was maintained along otherwise imbalanced X-chromosome activity.

As lineages are transcriptionally highly distinct in post-implantation embryos^23^ (E5.5-6.5) (**Fig. 2f**), we continued the analyses in a lineage-specific manner. Whereas extraembryonic lineages (visceral endoderm, VE; extraembryonic ectoderm, ExE) retain iXCI in female cells, EPI cells undergo XCR followed by rXCI (**Fig. 2c**). Strikingly, we found that cells of the extraembryonic lineages maintained XCU along with iXCI (**Fig. 2g**) whereas female EPI cells residing in reactivated XaXa state (XCR state) lacked XCU regardless of the embryonic age of the cells (**Fig. 2g and Extended Data Fig. 2d**), importantly demonstrating the erasure of XCU *in vivo*. This was followed by a second wave of XCU in cells in which rXCI was either underway or completed (**Fig. 2g and Extended Data Fig. 2d**), further confirming that XCU regulation is highly dynamic and quantifiable both *in vivo* and *in vitro*. To control for effects related to differentiation, we inferred pseudotime trajectories for each lineage (**Extended Data Fig. 2e, Methods**) and performed linear regression, which demonstrated minimal association with X-linked expression compared to allele usage (**Extended Data Fig. 2f-g**), thereby highlighting that the XCI and XCU processes can be separated from differentiation at single-cell resolution^17^.

Together, our data reveal that XCU is initiated in response to imbalanced dosage in a sex-specific and dosage-dependent manner. In males, XCU co-occur with ZGA completion while female XCU transpires in parallel with iXCI, followed by erasure and reestablishment in the epiblast along XCR and rXCI; and ablation of XCI by *Xist* knockout obstructs female XCU.

### A quantitative relationship between X-chromosome upregulation and X-inactivation

Throughout the early embryonic development, we had found that XCU has a remarkable flexibility in balancing X-linked expression along allelic imbalances and phases of XCI progression. However, it remained unclear whether XCU responded quantitatively to the lowering of expression from the Xi allele. If *trans-acting* factors are gradually shifted towards the Xa allele upon XCI, as hypothesized from our previous work^24,25^, XCU may not show a simple on/off pattern but would tune expression according to the cellular degree of XCI silencing.

To ensure that such tuning-like dynamics could be measured on the allele-level across different levels of allelic imbalance, we constructed and sequenced mock libraries of equimolar concentration but containing various spiked ratios of purified C57 and CAST RNA (**Methods**). This control experiment showed that allelic ratios could be faithfully captured down to around 100,000 reads per sample or with as few as 100 genes using UMIs as well as reads (**Extended Data Fig. 3a-b**), signifying that the Smart-seq-based technique was well within the sensitivity range to detect tuning-like allelic regulation.

As rXCI is an asynchronous process^17,23^, it represents an ideal system to evaluate the XCU modus. Remarkably, as rXCI was established in EPI cells, the other allele displayed proportional chromosome-wide compensation (adjusted R^2^_maternal_ = 0.80, P_maternal_ = 2.92 × 10^−69^, adjusted R^2^_paternal_ = 0.91, P_paternal_ = 1.19 × 10^−103^, linear regression; **Fig. 3a**), in accordance with the tuning model. To further validate this tuning-like behavior we reanalyzed independent UMI-count data of embryonic stem cell priming generated by Pacini *et al*.^21^, indeed confirming our model (adjusted R^2^_CSR_ = 0.84, adjusted R^2^_CAST_ = 0.86, P < 2.2 × 10^−16^, linear regression; **Fig. 3b**). Strikingly, variability of iXCI completeness in extraembryonic lineages and pre-implantation stages also followed the trend projected from EPI cells (**Fig. 3a and Extended Data Fig. 3c**), suggesting that the first, iXCI-associated, wave of XCU is achieved by the same mechanism as that during rXCI.

**Figure 3.**
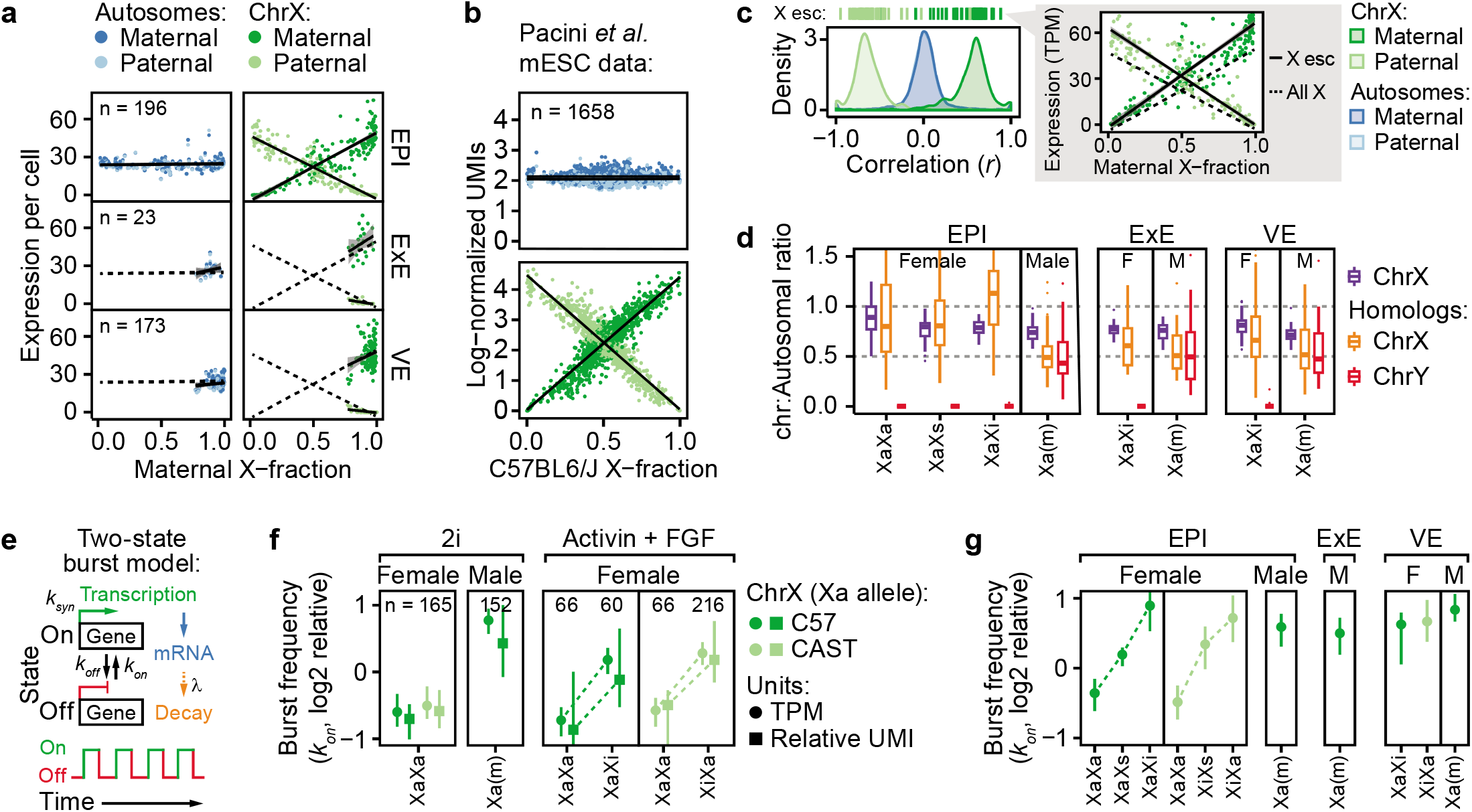
X-upregulation follows a tuning-like mode of regulation. **a.** Scatter of expression level per allele (y-axis) and maternal X-fraction per female cell (x-axis) with lines indicating linear model mean ± 95% confidence interval. Data shown for three lineages (EPI, ExE and VE). Dashed black lines project EPI expression. Shown for autosomes (blue; n = 14,941–17,643 genes) and chrX (green; n = 597–713). **b.** Same as panel a. but shown for mESCs cultured under serum/LIF conditions for up to 4 days (Pacini et al. 2021). Shown for autosomes (blue; n = 1,074–4,913 genes) and chrX (green; n = 23–182). Legend as in panel a. but for C57 and CAST alleles instead of maternal and paternal, respectively. **c.** Density plot of gene-wise correlations (Pearson) against maternal X-fractions in EPI cells. Inset shows XCI escapee (“X esc”) expression per cell and allele. **d.** X:Autosomal ratios per cell shown as box plots grouped by lineage, sex, and XCI state for chrX (n = 129–410 genes) and ancestral X-Y homologs (n = 5–9 genes) separately. **e.** Inference of transcriptional kinetics from a two-state model of transcription (see Larsson et al. 2019 for details). **f.** Transcriptional burst frequency (*k_on_*) for active X alleles (Xa per parental strain) grouped by sex, culture condition and XCI state, shown as median ± 95% confidence interval, inferred by either TPM (dot) or relative UMIs (square). Data shown relative to median autosomal burst frequency. n genes = 10,836–16,714 and 348–545 for autosomes and chrX, respectively. **g.** Transcriptional burst frequency (*k_on_*) of the Xa allele in cells of different lineage, sex, and X-inactivation state in mouse embryogenesis *in vivo*, shown as median ± 95% confidence interval relative to median autosomal burst frequency, n = 252–316 genes. Legend as in panel f., female ExE lineage lacked sufficient cells (n ≤ 20) for kinetic inference.

Together, these findings provide evidence for continuous feedback of the X alleles in accordance with the tuning model of XCU. Surprisingly, genes known to escape XCI showed similar trends as other X-linked genes in female cells (P_maternal_ = 0.62, P_paternal_ = 0.71, Kolmogorov–Smirnov test; **Fig. 3c**), which would be unlikely if X-linked expression is independently regulated per gene without a higher-level allelic logic. The notion that XCI escapees are subject to dosage balancing would help to explain why these genes are generally expressed at lower levels from the inactive X allele^26–28^. A subset of escapee genes have ancestral homologs remaining on the Y chromosome^29^. Remarkably, these X-Y gene pairs were not dosage compensated in males whereas the corresponding X-linked homologs displayed XCU in female cells of all lineages (Males: P > 0.05, Females: P_EPI_ = 6.02 × 10^−30^, P_VE_ = 3.91×10^−15^, P_ExE_ = 4.84×10^−2^, one-sample Wilcoxon test, μ = 0.5; **Fig. 3d**), further suggesting an allelic dosage balancing circuitry that goes beyond individual genes. Thus, the combined effect of biallelic expression and gene-specific XCU may explain why certain escapee genes tends to be expressed at higher levels in females^1^.

We hypothesized that the reason why the tuning-like mode of X-linked expression regulation was overlooked in previous studies was due to the lack of allele-resolution measurements of dosage compensation^8–12,14,21,30^. Indeed, all approaches we tested utilizing total expression levels (total X expression, X:Autosomal ratios, Female:Male ratios) failed to identify tuning-like XCU as all non-XaXa cell states produced similarly balanced expression (**Extended Data Fig. 3d-f**), explicitly exposing the risk of inferring allelic processes from non-allelic measurements.

Steady-state mRNA levels are determined by synthesis and degradation, suggesting that at least one of the two is altered to achieve the tuning-like effect of XCU. We have previously shown that expression-matched X-linked- and autosomal transcripts have similar decay rates^25^ whereas others have found XCU in steady state to be associated with increased transcriptional initiation^11,12,31^, suggesting that XCU is primarily controlled by transcriptional rather than post-transcriptional means. Furthermore, we dissected expression levels into kinetic parameters of transcriptional bursting (assuming a two-state model of transcription [on/off]; rate of transcription, burst frequency *k_on_* and burst size [*k_syn_*/*k_off_*]; **Fig 3e**), revealing that XCU is driven by increased transcriptional burst frequency^25,32^. To test whether bursting patterns reflect the tuning-like mode of XCU, we inferred transcriptional kinetics from our *in vitro* mESC model at the allelic level using both molecule-(UMI) and read-count (TPM) Smart-seq3 data (**Methods**). Indeed, the two XaXa alleles displayed moderate and balanced burst frequency (*k_on_*) whereas all states of monoallelic X-chromosome expression (*i.e*. XCU states) displayed markedly increased burst frequency (FDR_TPM_ ≤ 3.97 × 10^−10^, FDR_UMI_ ≤ 5.68 × 10^−03^, FDR-corrected Mann-Whitney *U* test; **Fig. 3f**) whereas burst size (*k_syn_*/*k_off_*) remained largely unchanged (FDR ≥ 0.08, **Extended Data Fig. 3g**), consistent with our previous report^25^. If XCU acts through tuning-like dynamics, we expected a gradual increase in burst frequency from the Xa allele as XCI progressed on the other. To test this, we inferred transcriptional kinetics using the *in vivo* scRNA-seq data and indeed found that burst frequency on the Xa allele was progressively increased during rXCI establishment in epiblasts (**Fig. 3g**, **Extended Data Fig. 3h**). Interestingly, burst frequency was lost at the Xi allele at a higher rate than it was gained at the Xa allele (**Extended Data Fig. 3i**), suggesting that residual mRNA molecules from the silenced allele help buffer the allelic balance. We also investigated the transcriptional kinetics in extraembryonic cells subject to iXCI (**Fig. 3g**) indeed observing elevated burst frequency on the hyperactive X allele, pointing towards a general mechanism of allelic tuning of XCU by transcriptional bursting.

Together, we show that XCU acts in a tuning-like mode of action to dynamically achieve balanced RNA dosage via modulation of transcriptional burst frequency on the active X allele in mESCs priming *in vitro* as well as early embryogenesis *in vivo*. Furthermore, we extend this concept in a gene-wise manner by revealing that XCI escapees and X-Y homologs follow the same dosage balancing principles as other X-linked genes, suggesting a unified mechanism across genes on the X chromosome.

### Chromatin features of the X-chromosome upregulation state

Transcriptional burst frequency and size are primarily regulated by enhancer and promoter elements, respectively^32^, suggesting that enhancer activity may help drive XCU. A previous study reported increased histone acetylation levels upon XCU relative to autosomes^12^, begging the question whether X-linked enhancers reside in a more accessible state during hyperactivation.

To address the regulatory state of XCU, we developed a single-cell multi-modal profiling assay combining Smart-seq3 and scATAC-seq in parallel in the very same cells (**Methods**). We applied this method to our mESC priming model throughout rXCI and XCU establishment (**Fig. 4a, Extended Data Fig. 4a**), providing the first combined scRNA/ATAC-seq profiling with allelic resolution to our knowledge. As expected, the mESC-to-EpiSC transition was accompanied by distinct changes in chromatin accessibility of pluripotency and differentiation markers (**Fig. 4b-c**), matching differential expression in our initial Smart-seq3-only experiment (Odds ratio = 2.66, P = 8.34 × 10^−15^, Fisher’s Exact Test). Next, we integrated single-cell expression and chromatin accessibility measurements, which further confirmed the agreement between the RNA and DNA modalities (**Fig. 4d, Extended Data Fig. 4b**). To assess whether our combined scRNA/ATAC-seq assay could accurately detect gene- and allele-specific accessibility, we investigated imprinted autosomal genes^33^ which showed expected allelic skewing in both expression and accessibility (**Fig. 4e-f**), signifying sufficient sensitivity to detect allele-specific effects. Next, we grouped cells into different XCI states based on allelic expression in the RNA modality, which demonstrated concurrent loss of chromatin accessibility on the Xi allele (**Fig. 4g**) whereas autosomes remained biallelically accessible (**Extended Data Fig. 4c**). Surprisingly, unlike the distinct shift in X-linked RNA levels, accessibility of the active allele did not increase upon XCU neither in male nor female cells (**Extended Data Fig. 4d**). This was also the case when matching expression and accessibility for the same gene where only RNA-level upregulation was observed relative to the biallelic XaXa state, regardless of the degree of XCU (**Extended Data Fig. 4e-f**). This indicates that the transcriptional action of XCU transpires on a basal state of chromatin openness, making it unlikely that the lower allelic expression levels in the XaXa state is due to partial repression. To explore potential spatial patterns of XCU, we separated alleles based on degree of XCI completion which indeed revealed unchanged accessibility across the Xa allele whereas Xi accessibility was preferentially lost at regions gaining the heterochromatin histone modification H3K27me3 (**Fig. 4h**).

**Figure 4.**
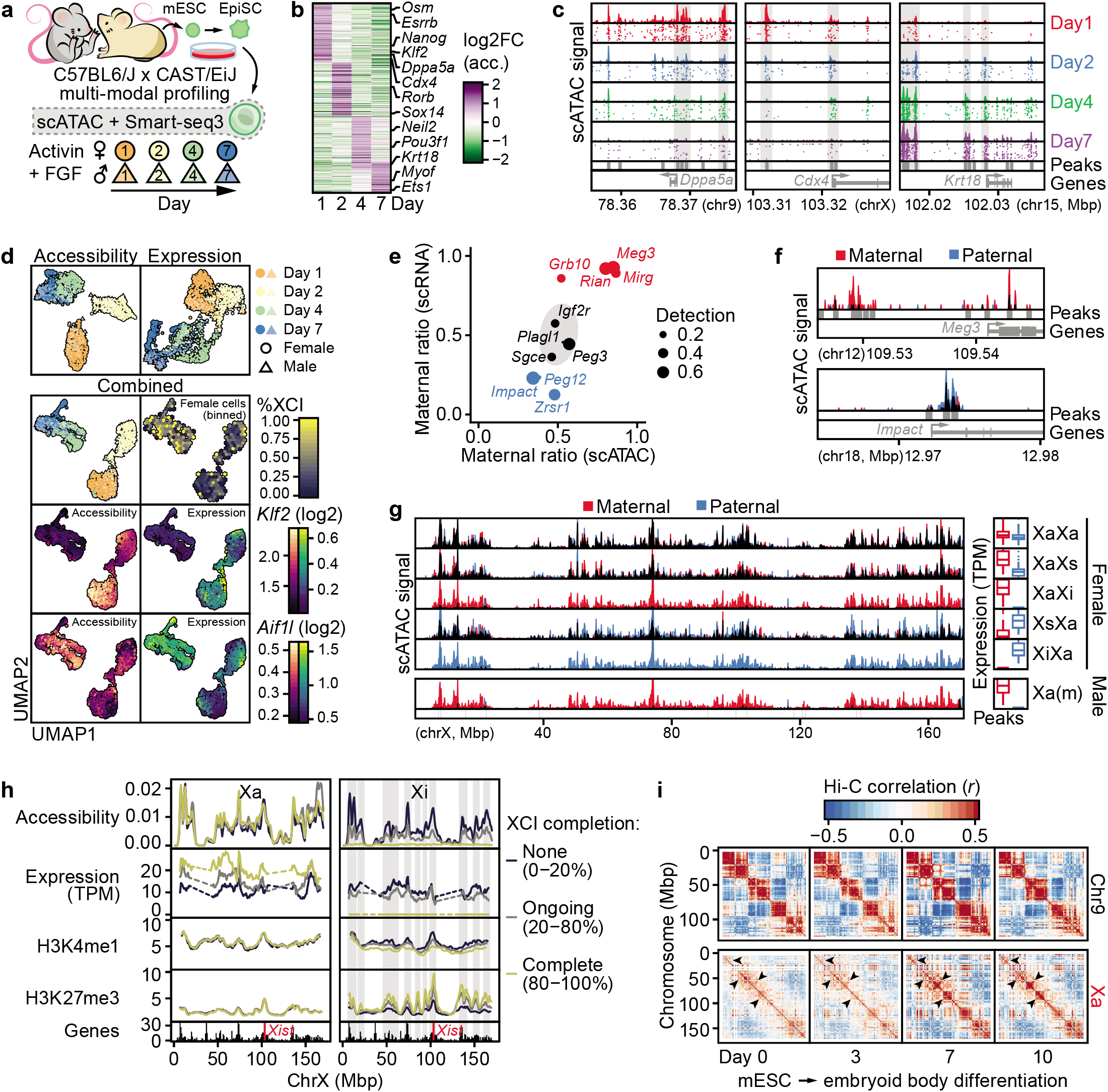
Epigenetic state of the hyperactive X chromosome. **a.** Schematic overview of experimental setup. C57BL6/J × CAST/EiJ F1 hybrid male and female mESCs were primed to EpiSCs using Activin A and FGF2 for up to 7 days to induce X-inactivation (XCI). Cells were collected at conditions and days of priming as indicated and subjected to allelic single-cell multi-modal profiling of expression (Smart-seq3) and chromatin accessibility (scATAC). **b.** Heatmap of accessibility changes along EpiSC priming. Acc. = accessibility gene score. **c.** Genome tracks of representative genes changing accessibility along EpiSC priming. Shown as single-cell accessibility and merged pseudobulk tracks. **d.** UMAP dimensionality reductions for accessibility (top 25,000 features), expression (top 1,000 HVGs), or the combined dimensions per cell. **e.** Maternal ratios for imprinted genes based in scRNA-seq (y-axis) and scATAC-seq (x-axis). Detection = detected in fraction of cells. **f.** Genomic tracks of allelic pseudobulk accessibility for representative genes from e. **g.** Genomic tracks of allelic pseudobulk accessibility for the entire X chromosome. Cells grouped based on XCI state and sex with corresponding expression of chrX shown to the right. **h.** Rolling average across each X allele, grouped based on degree of XCI completion. Allelic native ChIP-seq (Zylicz et al. 2019) shown for H3K4me1 and H3K27me3 as none, ongoing, or complete XCI for 0, 12 and 24h or DOX-induced XCI, respectively. **i.** *In situ* Hi-C contact correlation maps shown for the active X allele (Xa) and chr9 (CAST allele, corresponding to Xa) along mESC differentiation.

The observed lack of increased chromatin accessibility in the XCU state was surprising as the X chromosome has previously been associated with increased levels of permissive histone modifications, including acetylation, relative to autosomes^12,31^. However, as this was primarily studied in differentiated cells in steady-state XCU, and the measure normalized to autosomes, the effect may represent an inherent feature of the X chromosome rather than related to XCU. To address this further, we reanalyzed allelic native ChIP-seq (H2AK119Ub, H3K27me3, H3K4me1, H3K4me3, H3K27ac, H3K9ac, H4ac) time-series data for mESCs with DOX-inducible XCI^34^. In agreement with our combined scRNA/ATAC-seq data, the number of enriched regions for all histone modifications remained constant on the Xa allele throughout XCI/XCU progression (**Extended Data Fig. 4g**), as did modification density at both promoters and enhancers (**Extended Data Fig. 4h**). This corroborates our initial finding that the chromatin accessibility landscape remains unchanged upon XCU.

Intrigued by these observations at the chromatin level, we hypothesized that the burst-frequency-driven XCU may not be modulated by enhancer activity *per se* but through enhancer-gene contacts^35,36^. To explore this, we analyzed allele-resolved timeseries *in situ* Hi-C data for mESC undergoing differentiation for up to 10 days^37^(**Methods**). Unlike the Xi allele, that assumes a distinct bipartite mega-domain structure upon XCI^37–39^, Xa retained its global long-range chromosome conformation (**Extended Data Fig. 5a**). However, the shorter-range chromatin domains on Xa became increasingly distinct as the cells underwent XCU (**Fig. 4i, Extended Data Fig. 5b-c**), suggesting that XCU is associated with increased chromatin contacts, consistent with the observed increase in transcriptional burst frequency. Interestingly, a recent experimental study of XCR also captured increased compartmentalization of the Xa allele in differentiated cells and intermediately reprogrammed induced pluripotent stem cells (iPSCs) relative to mESCs^40^, but the connection to XCU was not made in the study as the XCU dynamics during reactivation was first pinpointed in the present study.

Together, we performed epigenetic profiling of XCU establishment, revealing that chromatin accessibility and histone modification H2AK119Ub, H3K27me3, H3K4me1, H3K4me3, H3K27ac, H3K9ac, and H4ac remain in a basal state upon XCU whereas shorter-range chromatin contacts and compartmentalization are increased, suggesting that enhancer-gene contacts help drive its tuning-like modulation of burst frequency.

## Discussion

In this study, we have uncovered an unanticipated flexibility of mammalian XCU in controlling X-linked expression, with fundamental implications to dosage compensation. By tracing expression levels at allelic resolution in single cells during early murine embryonic development, we identify key sex-specific events and timing for initiation, maintenance, erasure, and re-establishment of allelic X-upregulation to compensate otherwise unbalanced RNA levels. Notably, we show that XCU is achieved by transcriptional burst frequency increase as a universal kinetic drive, across sexes, cell lineages, *in vivo* and in experimentally controlled systems, co-occurring with increased chromatin compartmentalization. In contrast to XCI, which is initiated in a tightly controlled manner^22^, we find that XCU acts in a flexible tuning-like fashion. We further demonstrate that XCI is required to initiate XCU in female (XX) cells, indicating that XCU occurs as a direct response to imbalanced X dosage. This is distinct from previous reports suggesting XCU to be established already in the zygote^8^ or progressively after the 4-cell stage in both sexes^14,15^. These discrepancies may be explained by the inability of non-allelic gene-expression measurements to correctly distinguish XCU in the presence of parallel confounding allelic processes such, as ZGA and XCI, whereas our present study directly attributes XCU to the active X allele. Our surprising finding that female cells with biallelic XaXa expression (naïve female mESCs, 4-cell embryos and epiblasts) lack previously assumed XCU^10,11,41^ is in line with observations of balanced X:A dosage in haploid cells (including MII oocytes; **Fig. 2d**) or other cellular states harboring two active X chromosomes, such as primary oocytes and primordial germ cells^8,30,42,43^. Interestingly, the higher total dosage of the XaXa state that we uncovered may also explain a recent report showing higher X:A dosage in human female primordial germ cells compared to males^44^, suggesting that our findings in mouse translate to humans. Furthermore, the gene expression dosage of two Xa alleles is known to be incompatible with sustained embryonic development^22,45^ which fits with the observation of incomplete compensation by XCU at the RNA level (~1.6 fold) reported by us and others^8–12,25,31^ (**Fig. 1**). Using allele-resolved multi-omics, including a novel joint scRNA/ATAC-seq assay, we characterized the epigenetic landscape of XCU. In contrast to readily observable differentiation- and XCI-related changes, we found the chromatin landscape of XCU to be unaltered while shorter-range DNA contacts and compartmentalization increased. The notion that XCU is driven by increased transcriptional burst frequency and chromatin contacts may explain why active histone modifications do not scale proportionally with RNA levels on the hyperactive X allele^12,31^.

Based on our data, we provide a revised model of the sequence of events by which mammalian X-chromosome dosage compensation is achieved in the presence of other allelic processes during development (*i.e*. ZGA, iXCI, XCR, and rXCI), schematically summarized in **Fig. 5**. This alters the widely-held notion that XCI acts on hyperactive Xa alleles in females^10,11,41^ (‘Default XCU’ model; **Fig. 5a top**). Instead, our data promotes a model where the two Xa alleles are moderately expressed and that XCU gradually tunes Xa expression levels throughout XCI proportional to dosage imbalances (‘Dynamic XCU’ model; **Fig. 5a middle**). As dosage compensation through XCU is less than 2-fold, the lethal dose of two hyperactive X alleles is avoided in the ‘Dynamic XCU’ model (**Fig. 5a bottom**). It is important to note that the transcription-driven XCU we observe may act on top of other layers of regulation at post-transcriptional- and translational levels^1^. Our findings suggest that increased transcription factor concentrations at the hyperactive Xa allele play a key role in its regulation. Not only is the single Xa allele subjected to higher doses of factors transcribed from diploid autosomes, but the increased chromatin contacts we observe concurrent with XCU (**Fig. 4i**) may increase local transcription factor concentrations through loop-mediated trapping^46^. As transcriptional burst frequency is controlled by both local factor concentrations^47^ and enhancer-promoter contacts^32,35,36^, this would ultimately result in the increased transcriptional burst frequency in XCU we observe (**Fig. 3f-g**). Furthermore, the same model can also operate for XCI in females as transcription factors are rapidly excluded from the Xi allele when repressive compartments are established^48^. As XCI is a gradual process^19^, factor complexes could progressively shift to the Xa allele in line with the tuning-like mode of XCU. Collectively, this would result in balanced RNA levels, explaining how the total dosage of X-encoded transcripts are maintained so surprisingly stable throughout development and XCI progression in mammalian species^8,10,13,14,49^ and why XCI escapees are expressed at higher levels from the Xa allele^26–28^ whereas X-Y homologs are not dosage compensated in males as both chromosomes are active (**Fig. 3d**). Finally, as two active X chromosomes is considered a hallmark of the naïve female stem cell state and a gold standard in reprogramming studies^50^, our finding that the biallelic XaXa state lacks XCU alters the interpretation of X-chromosome expression level measurements for assessment of reprogramming success and naïveness.

**Figure 5.**
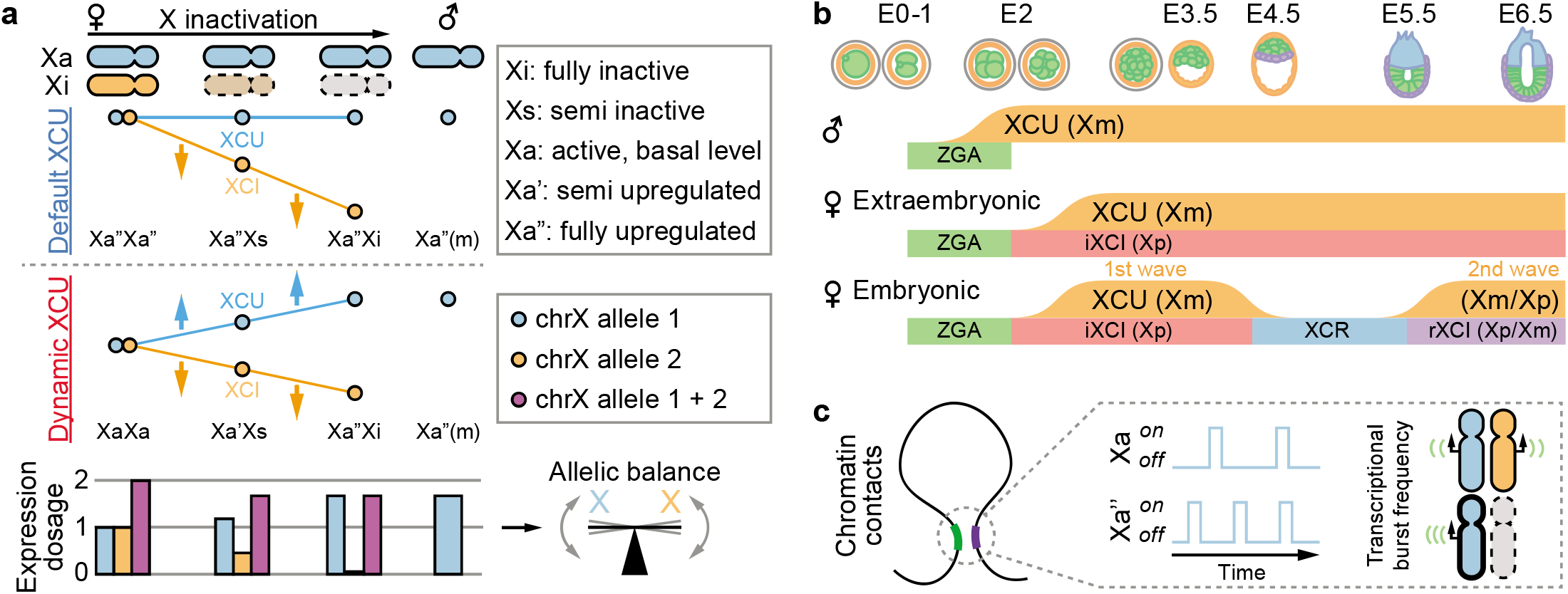
A unified model of dosage compensation by X-upregulation and inactivation. **a.** A previous model of X-upregulation (XCU) postulated that XCU was active in XaXa and all states of X-inactivation (XCI) [‘Default XCU’] in female cells, whereas our experimental allele-level data enforce an updated model in which XCU is tuned in relationship to the degree of XCI [‘Dynamic XCU’] (top). However, the single hyperactive allele (Xa”) does not fully reach the combined expression level of the two moderately active alleles in female XaXa state, and total output is kept allelically balanced in non-XaXa states (bottom). **b.** Overview of XCU timing dynamics across mouse pre- and early post-implantation development in the two sexes and embryonic/extraembryonic lineages. Abbreviations: zygotic genome activation (ZGA), imprinted X-chromosome inactivation (iXCI), X-chromosome reactivation (XCR), random X-chromosome inactivation (rXCI). **c.** As one allele is gradually silenced, the concentration of available transcription factors is shifted from the inactive (Xi) towards the active allele (Xa), increasing transcriptional burst frequency and chromatin-contact frequency of the hyperactive X allele (Xa’’).

Thus, in summary, our study provides comprehensive characterization and mechanistic insight into the allelic regulation of the murine X chromosome, prompting a revised model of dosage compensation that unifies the temporal dynamics of X-inactivation and X-upregulation.

## Supporting information

Supplementary Table 1 (Microsoft Excel format)

Supplementary Table 2 (Microsoft Excel format)

## Data availability

Raw and pre-processed data generated is publicly available at ArrayExpress under accession E-MTAB-9324 (Smart-seq3) [*Reviewer access:* Username: Reviewer_E-MTAB-9324; Password: W98URLC1], E-MTAB-10709 (Allelic dilution series) [*Reviewer access:* Username: Reviewer_E-MTAB-10709; Password: kjdicxfA] and E-MTAB-10714 (Combined Smart-seq3+scATAC) [*Reviewer access:* Username: Reviewer_E-MTAB-10714; Password: qgieeder]. Previously published raw data is available at Gene Expression Omnibus under accessions GSE45719, GSE74155, GSE109071, GSE116480, GSE23943, GSE80810, GSE90516, GSE116649 and GSE151009.

## Code availability

Data and code generated during this study are available at github: github.com/reiniuslab/Lentini_XCU_in_vivo.

## Acknowledgements

This study was made possible by grants to from the Swedish Research Council (2017-01723), the Ragnar Söderberg Foundation (M16/17), and SRA Stem Cells and Regenerative Medicine (Karolinska Institutet) to BR. ME is supported by The Swedish Cancer Society, The Swedish Childhood Cancer Fund, Radiumhemmets forskningsfonder, SFO StratRegen, The Swedish Research Council (2020-02940) and Cancer Research KI. AL is supported by a postdoctoral fellowship from the Swedish Society for Medical Research. We thank members of the Reinius lab and Colm Nestor for comments on the manuscript, and Björn Högberg and Ana Teixeira for use of an Illumina sequencer.

## Contributions

A.L.: Investigation, Methodology, Formal analysis, Conceptualization, Visualization, Data Curation, Writing - Original Draft, Writing - Review & Editing. H.C.: Investigation. J.C.N.: Investigation. N.P.: Investigation. C.C.: Investigation. N.A.: Investigation. M.E.: Methodology, Resources. Q.D.: Methodology, Resources. B.R.: Investigation, Methodology, Conceptualization, Supervision, Resources, Project administration, Funding acquisition, Writing - Original Draft, Writing - Review & Editing.

**Corresponding authors**

Correspondence to

## Supplementary information

Materials and Methods

Extended Data Figures 1-5

Supplementary Tables 1-2

Supplementary References

---

**Extended Data Fig. 1.**
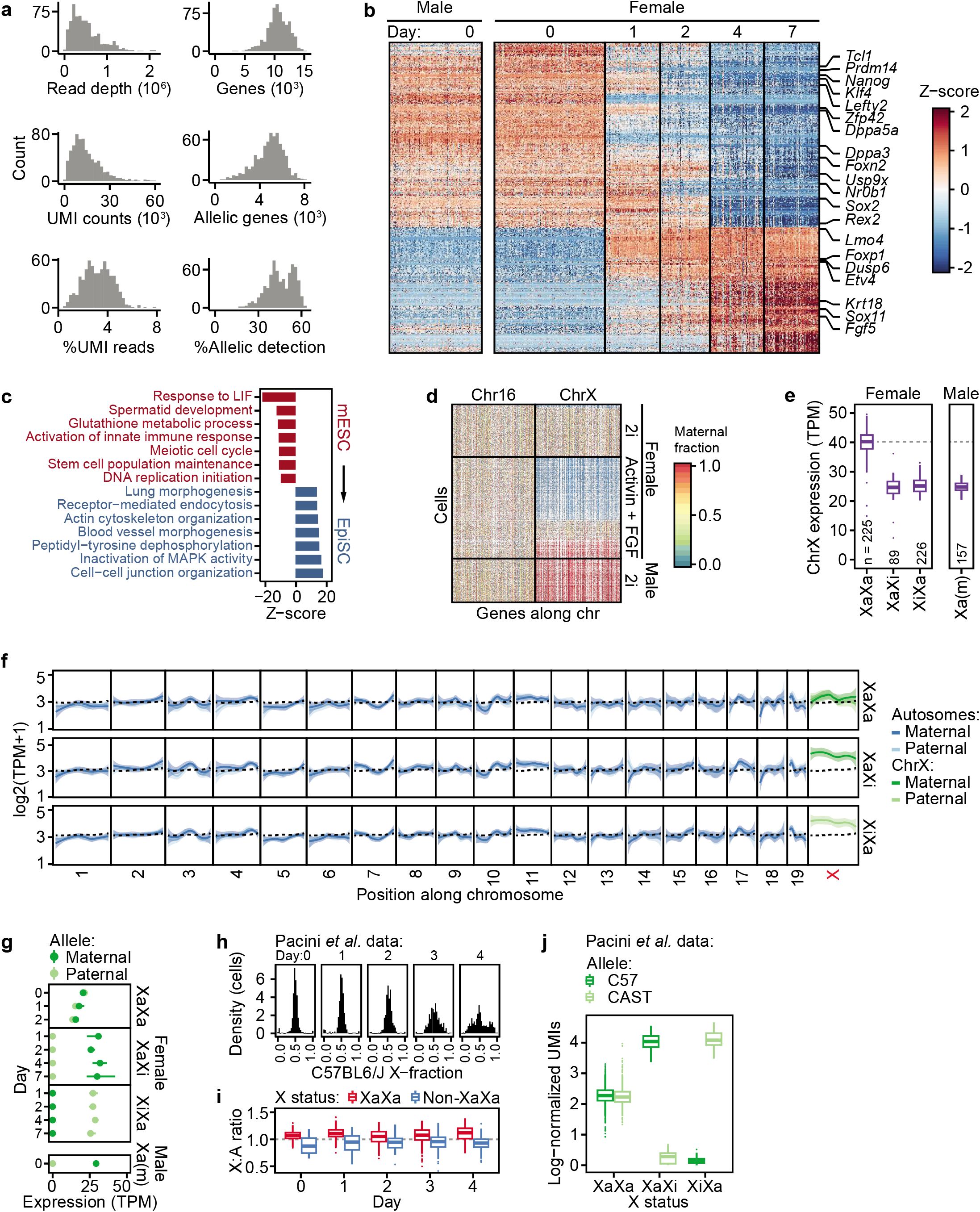
Validation of X-upregulation upon exit from pluripotency. **a.** Histograms of read and gene metrics for generated Smart-seq3 data. **b.** Heatmap of differentially expressed genes along mESC (day 0) priming towards EpiSCs (day 1-7). See also **Supplementary Table 1**. **c.** GO term enrichment for genes in **b.** See also **Supplementary Table 1**. **d.** Heatmap of maternal fractions for chr16 and chrX for genes detected in >10% of cells for mESCs under 2i or EpiSC priming conditions. **e.** Boxplot showing total expression (RNA output of both alleles) for X-linked genes (n = 1,158–1,337) grouped by XCI state. Dashed line indicates the XaXa median. **f.** Rolling average (LOESS fit ± 95% confidence interval) of allelic gene expression along intra-chromosomal coordinates (x-axis) for female mESCs cells cultured in EpiSC priming conditions grouped by XCI state. Dashed line indicates the average for autosomes. **g.** Allele-resolved chrX expression grouped by sex, XCI state and timepoints of EpiSC priming, shown as median ± 95% confidence interval. **h.** Density plots of allelic ratios for mESCs cultured under serum/LIF conditions for up to 4 days. **i.** Boxplots of X:Autosomal ratios for cells in **h**, grouped by XCI state per day. **j.** Boxplots of allelic expression for cells in **h-i**, grouped by XCI state.

**Extended Data Fig. 2.**
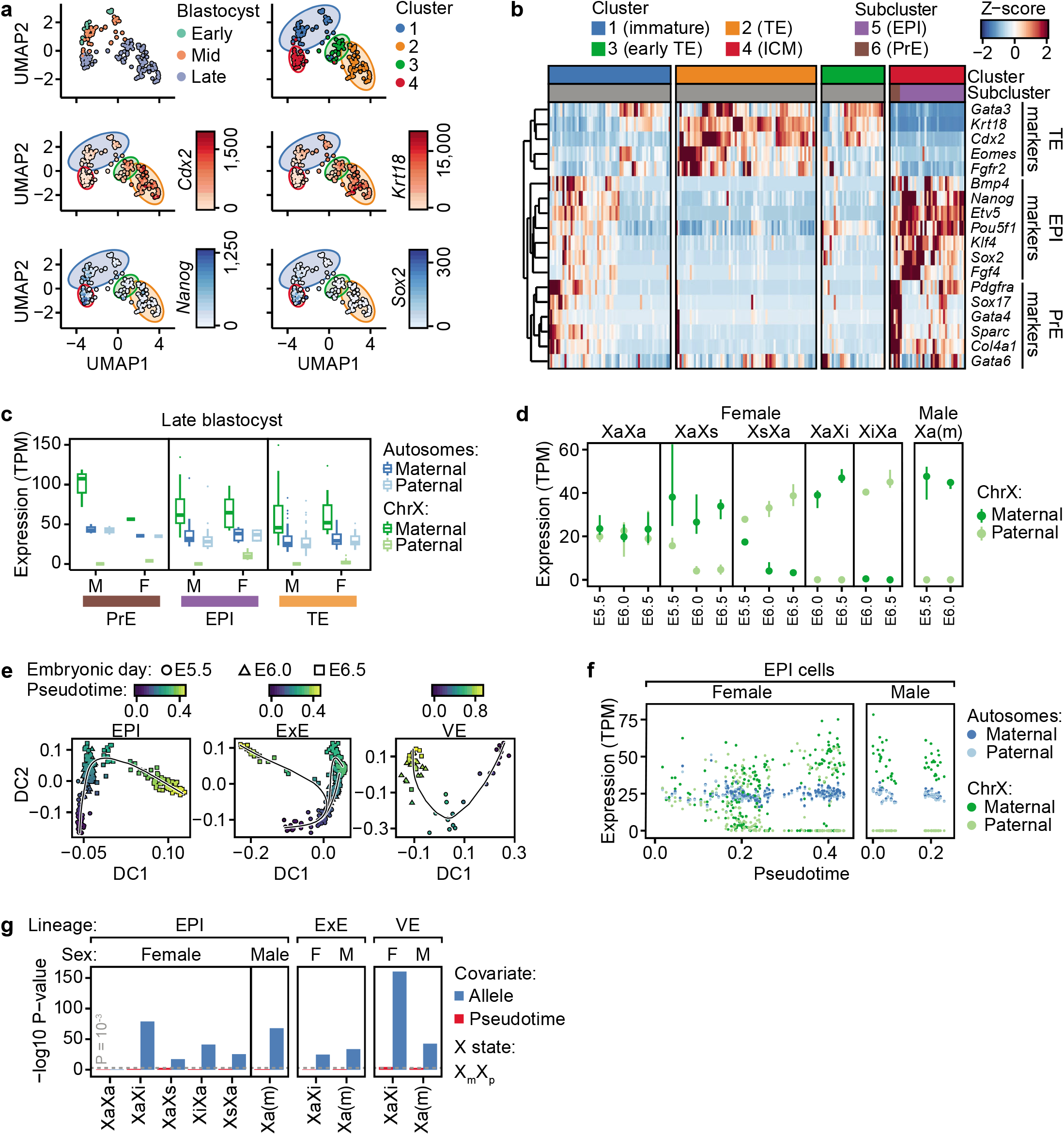
Extended analyses of early embryo development. **a.** UMAP dimensionality reduction for pre-implantation blastocysts using the top 1,000 variable genes and graph-based clustering of blastocysts. Trophectoderm-(TE) and inner cell mass (ICM) marker genes shown in red and blue, respectively. **b.** Heatmap and hierarchical clustering of pre-implantation blastocysts using lineage-specific marker genes for trophectoderm (TE), epiblast (EPI), primitive endoderm (PrE) lineages. **c.** Boxplots of allelic expression of late blastocyst subclusters identified in (**a-b**). **d.** Allelic ChrX expression of post-implantation epiblast (EPI) cells grouped by embryonic day and XCI state, shown as median ± 95% confidence interval. **e.** Diffusion map dimensionality reduction and Slingshot trajectory inference per post-implantation lineage. Embryonic day indicated as E5.5 (circle), E6.0 (triangle) or E6.5 (square). **f.** Allelic expression along pseudotime trajectories from **e** for post-implantation EPI cells. **g.** Association of allele usage or pseudotime on allelic expression using linear modelling.

**Extended Data Fig. 3.**
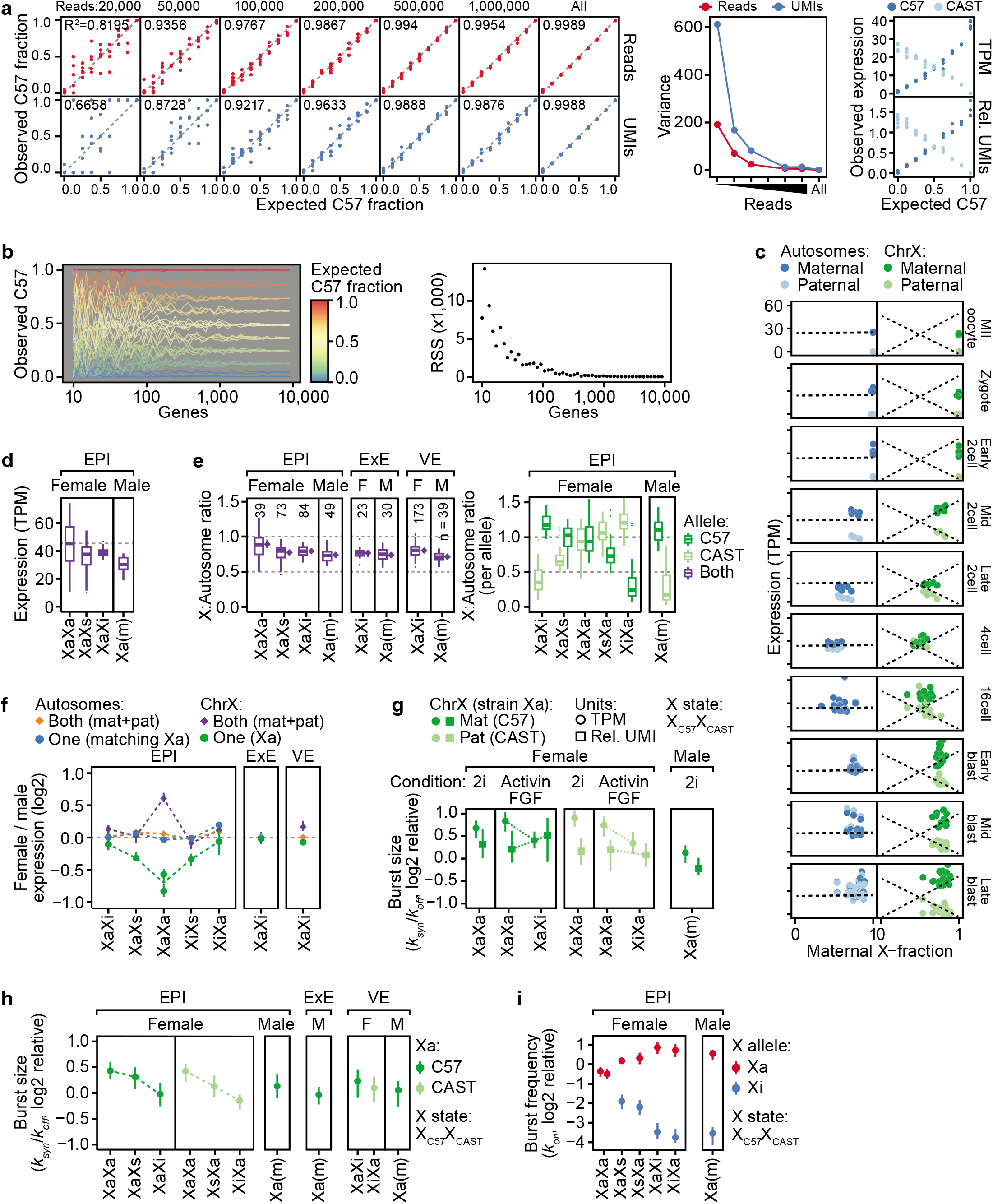
Extended analyses of relationship between X-upregulation and X-inactivation. **a.** Accuracy of allelic inference of Smart-seq3 using experimentally controlled ratios of C57:CAST RNA, based on read count down sampling. R^2^ and variance (σ^2^) calculated from linear model. **b.** Stability of allelic inference of Smart-seq3 based on gene subsampling. RSS = residual sum of squares. **c.** Relationship between X expression and maternal X-fraction in pre-implantation embryos, dashed lines represent post-implantation EPI trends. **d.** Boxplots of total chrX expression (accumulative RNA output of both alleles) in EPI cells grouped by XCI state. **e.** X:Autosomal ratios shown as box plots or bootstrapped median (diamond ± 95% confidence) interval for total expression (both alleles; left) or allelic resolution (right). Data stratified by lineage, sex and XCI state. F = female, M = male. **f.** Female:male expression ratios calculated either by total expression (accumulative from both alleles; diamond) or for one active X allele (Xa; dot), shown as median ± 95% confidence interval. For epiblasts (EPI) the data points are stratified according to female rXCI state (x-axis). Autosomal genes (n = 14,941–17,643). X-linked genes (n = 597–713) genes. **g.** Transcriptional burst size (*k_syn_*/*k_off_*) for active X alleles (Xa) grouped by sex, culture condition and XCI state, shown as median ± 95% confidence interval, inferred by either TPM (dot) or relative UMIs (square). The data is shown relative to median autosomal burst size. **h.** Transcriptional burst size (*k_syn_*/*k_off_*) for active X alleles (Xa) grouped by sex, lineage and rXCI status, shown as median ± 95% confidence interval. The data is shown relative to median autosomal burst size. Note however that the accuracy of burst-size inference is limited in non-UMI (Smart-seq2) scRNA-seq data (See Larsson et al. 2019 for details). **i.** Transcriptional burst frequency (*k_on_*) for Xa and Xi alleles in EPI cells grouped by sex and rXCI status, shown as median ± 95% confidence interval. The data is shown relative to median autosomal burst frequency.

**Extended Data Fig. 4.**
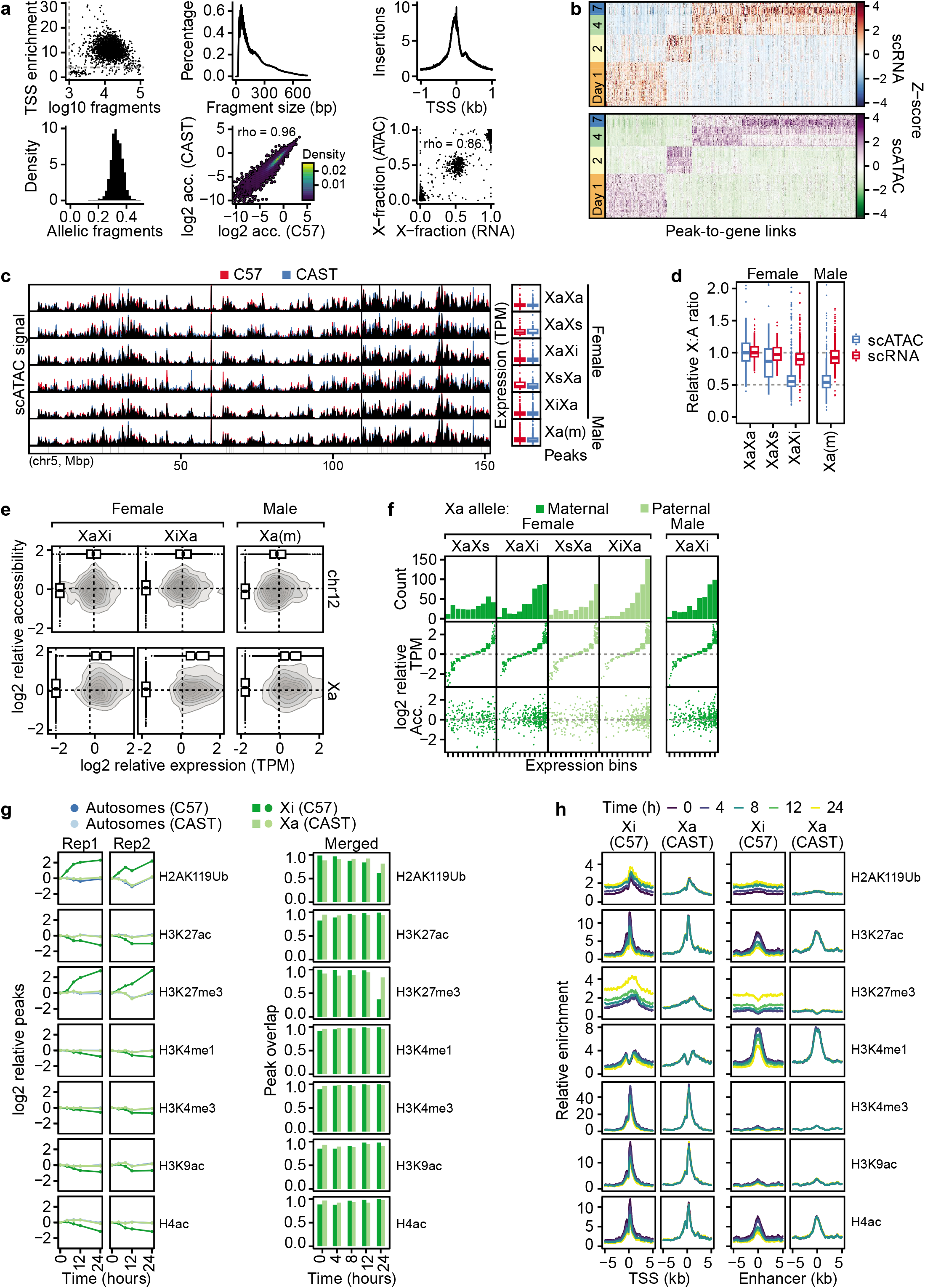
Extended analyses of genomic regulation of X-upregulation. **a.** Quality metrics for generated combined scRNA/ATAC-seq data. **b.** Heatmaps of linked accessibility and expression along EpiSC priming. **c.** Genome tracks of allelic accessibility for an autosome (chr5), grouped by X-inactivation (XCI) state. Corresponding allele-resolved expression is shown to the right. **d.** Boxplots of X:Autosomal ratios relative to XaXa cells for expression (scRNA) and accessibility (scATAC). **e.** 2D density plots of gene-level accessibility (y-axis) and expression (x-axis) relative to XaXa cells shown per sex and XCI state for the active X allele (Xa) and chr12 (allele corresponding to Xa). Dashed line indicates autosomal median per group. **f.** Same data as in **e** but shown as jitter plots binned by relative expression. **g.** Number of native ChIP-seq peaks per modification and allele relative to the 0h timepoint (left) and peak overlap with all other timepoints per timepoint (right). **h.** Normalized allelic enrichment profiles of histone modifications for Xa and Xi alleles shown relative to transcription start site (TSS; left) or enhancers (right).

**Extended Data Fig. 5.**
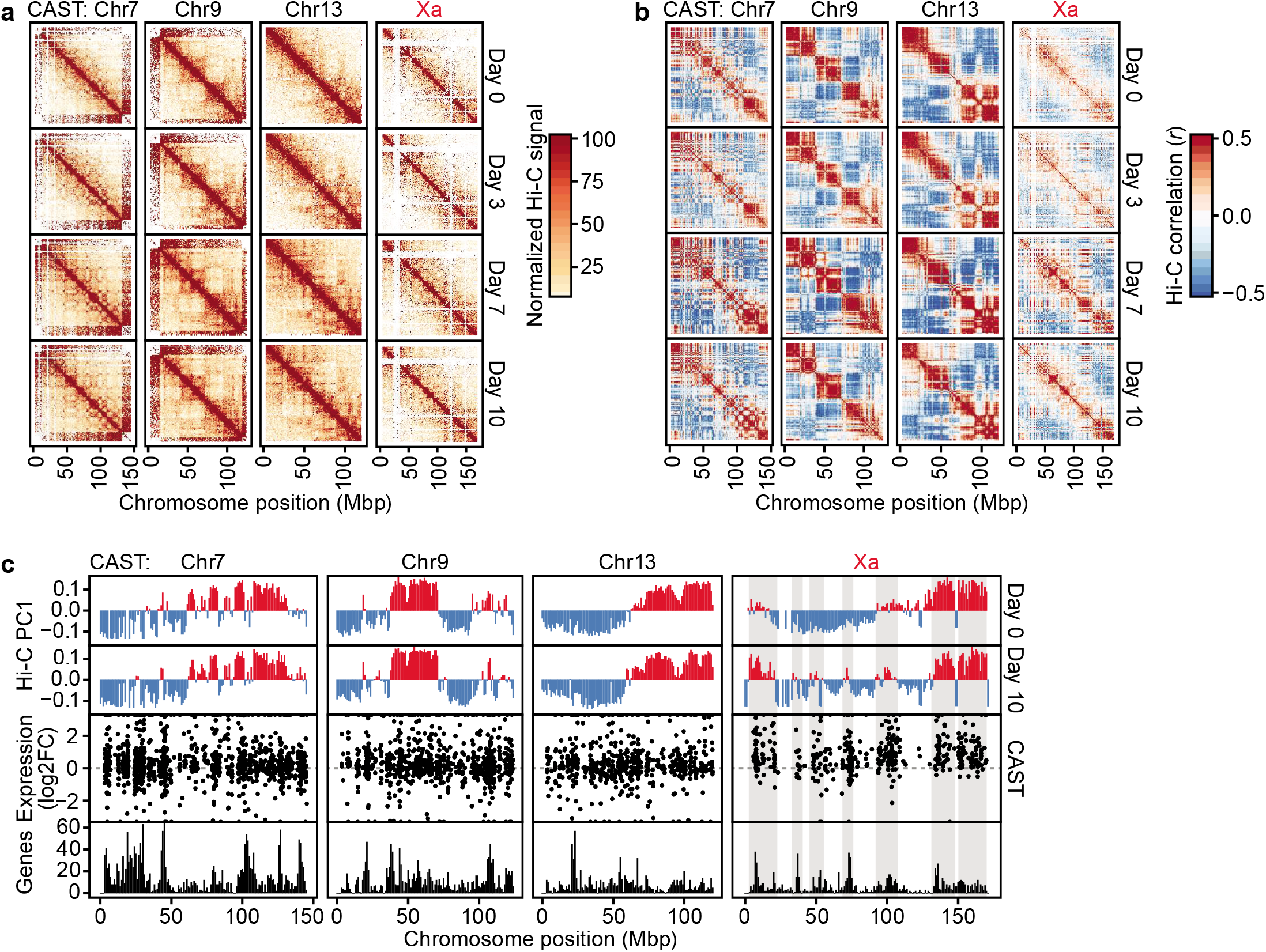
Extended analyses of chromatin contacts. **a.** Normalized *in situ* Hi-C contact signal maps for Xa and representative autosomes along mESC differentiation at 1Mb resolution. **b.** Same as **a** but shown as contact correlation maps. **c.** Eigenvector plots of chromosome compartmentalization. Also shown is Smart-seq3 relative expression of the CAST allele (XiXa *vs*. XaXa).

## SUPPLEMENTARY INFORMATION

### Materials and Methods

#### Ethics statement

All animal experimental procedures were performed in accordance with Karolinska Institutet’s guidelines and approved by the Swedish Board of Agriculture (permits 17956-2018 and 18729-2019 Jordbruksverket).

#### Derivation and culturing of cell lines

Male and female cell lines were established as previously described^1^. In brief, mESCs were derived from E4 blastocysts of F1 embryos (female C57BL/6J × male CAST/EiJ) and adapted to 2i condition by growing them in gelatin-coated flasks in N2B27 medium (50% neurobasal medium [Gibco], 50% DMEM/F12 [Gibco], 2 mM L-glutamine [Gibco], 0.1 mM β-mercaptoethanol, NDiff Neuro-2 supplement [Millipore], B27 serum-free supplement [Gibco]) supplemented with 1,000 units/mL LIF, 3 μM Gsk3 inhibitor CT-99021, 1 μM MEK inhibitor PD0325901, and passaged with accutase [Gibco]. To induce differentiation toward EpiSCs, mESCs grown in serum/LIF were plated on FBS coated tissue culture plates (coated overnight at 37°C) in N2B27 medium supplemented with 8 ng/mL Fgf2 (R&D) and 20 ng/mL Activin A (R&D) at a cell density of 1 × 10^4^ cells/cm^2^ and cultured for up to 7 days. Cells were split and re-plated in the same condition after 1, 2, 4 and 7 days of differentiation. At each split an aliquot of cells was collected for single-cell sorting into 96-well plates containing Smart-seq3 lysis buffer.

#### Single-cell RNA sequencing (Smart-seq3)

scRNA-seq libraries were constructed as previously described^2^ with slight modification. Briefly, cells were singlecell sorted into 96-well low-bind PCR-plates [Eppendorf] containing 3 μl of lysis buffer (0.5 units/μl RNase inhibitor [Takara], 0.15% Triton X-100 [Sigma], 0.5 mM (each) dNTP [Thermo Scientific], 1 μM oligo-dT primer [5′-biotin-ACGAGCATCAGCAGCATACGAT30VN-3′; IDT], 5% PEG [Sigma]). Sorting was performed using an SH800 [Sony]. Plates were briefly centrifuged immediately after sorting, sealed, and stored at −80°C. For cell lysis and RNA denaturation, plates were incubated at 72°C for 10 min and immediately placed on ice. Next, 5 μl of reverse transcription mix (50 mM Tris-HCl, pH 8.3 [Sigma], 75 mM NaCl [Ambion], 1 mM GTP [Thermo Scientific], 3 mM MgCl_2_ [Ambion], 10 mM DTT [Thermo Scientific], 1 units/μl RNase inhibitor [Takara], 2 μM of template-switch oligo [5′-biotin-AGAGACAGATTGCGCAATGNNNNNNNNrGrGrG-3′; IDT] and 2 U/μl of Maxima H-minus reverse transcriptase [Thermo Scientific]) was added to each sample. Reverse transcription was carried out at 42°C for 90 min followed by 10 cycles of 50°C for 2 min and 42°C for 2 min and the reaction was terminated at 85°C for 5 min. PCR pre-amplification was performed directly after reverse transcription by adding 17 μl of PCR mix (containing DNA polymerase, forward and reverse primer) bringing the final concentration in the 25 μl reaction to 1x KAPA HiFi ReadyMix [Roche] 0.1 μM forward primer [5′-TCGTCGGCAGCGTCAGATGTGTATAAGAGACAGATTGCGCAATG-3′; IDT] and 0.1 μM reverse primer [5’-ACGAGCATCAGCAGCATACGA-3′; IDT]). Thermocycling was performed as follows: 3 min at 98°C, 22 cycles of 20 s at 98°C, 30 s at 65°C and 6 min at 72°C, and final elongation at 6 min at 72°C. After PCR preamplification, samples were purified with AMpure XP beads [Beckman Coulter] at volume ratio 0.8:1. Library size distributions were monitored using high-sensitivity DNA chips (Agilent Bioanalyzer 2100) and cDNA concentrations were quantified using the Quant-iT PicoGreen dsDNA Assay Kit [Thermo Scientific]. cDNA was subsequently diluted to 100–200 pg/μl.

Tagmentation was performed using in-house produced Tn5^3^. 2 ng of cDNA in 5 μl water was mixed with 15 μl tagmentation mix (0.2 μl Tn5, 2 μl 10x TAPS MgCl_2_ Tagmentation buffer; 5 μl 40% PEG-8000; 7.8 μl water, per reaction) and incubated 8 min at 55°C in a thermal cycler. Tn5 was inactivated and released from the DNA by the addition of 4 μl 0.2% SDS and 5 min incubation at room temperature. Library amplification was performed by adding 5 μl mix of 1 μM of forward and reverse custom-designed Nextera index primers [forward: 5′-CAAGCAGAAGACGGCATACGAGATNNNNNNNNNNGTCTCGTGGGCTCGG-3′, reverse: 5′-AATGATACGGCGACCACCGAGATCTACACNNNNNNNNNNTCGTCGGCAGCGTCIDT-3’, where N represents the 10-bp index bases; IDT] and 15 μl PCR mix (1 μl KAPA HiFi DNA polymerase [Roche]; 10 μl 5x KAPA HiFi buffer; 1.5 μl 10 mM dNTPs; 3.5 μl H2O, per reaction), and thermal cycling: 3 min 72°C, 30 s 95°C, 13 cycles of 10 s 95°C; 30 s 55°C; 30 s 72°C, followed by final elongation at 5 min 72°C; 4°C hold. DNA sequencing libraries were purified using 0.8:1 volume of AMPure XP beads [Beckman Coulter]. Libraries were sequenced using a NextSeq 550 and High Output kits [Illumina].

#### Single-cell joint accessibility and RNA expression (scATAC+Smart-seq3)

The joint single-cell ATAC and SmartSeq3 analysis was performed as in DNTR-seq^4^, with the following modifications. Single cells were FACS sorted into 384 well plates containing 3μl lysis buffer (0.03ul 1M TrispH7.4, 0.0078 μl 5 M NaCl, 0.075 μl 10% IGEPAL, 0.075 μl RNase Inhibitor, 0.075 μl 1:1.2M ERCC, 2.7372 μl H2O). Immediately after sorting, plates were centrifuged at 1800 x g 4°C 5mins, placed on ice 5 mins, vortexed 3000 RPM 3mins, and centrifuged again at 1800x g 4°C 5mins to lyse the cells and spin down the nucleus. 2 μl of the supernatant was then carefully moved to a new 384 well plate for Smart-seq3 mRNA library preparation and the nucleus remained in the original well for scATAC library preparation. The scATAC *in situ* tagmentation was performed with 2 μl of Tn5 tagmentation mix (0.06 μl 1M Tris-pH 8.0, 0.0405 μl 1M MgCl_2_, 2 μl Tn5) and incubating at 37°C for 30 mins. After the tagmentation, 2 μl of the supernatant was aspirated and the nuclei were then washed once with 10 μl ice-cold washing buffer (0.1 μl 1M Tris-pH7.4, 0.02μl 5M NaCl, 0.03 μl 1M MgCl_2_, 9.85 μl H_2_O). The remaining Tn5 was inactivated by addition of 2μl 0.2% SDS-Triton X-100 followed by incubation at room temperature for 15 mins and 55°C for 10 mins. Barcoding PCR was done by KAPA HiFi PCR Kit [Roche] in a final 25μl reaction (11.5μl H_2_O, 5 μl 5X reaction buffer, 0.75 μl dNTP, 0.5 μl KAPA HiFi DNA Polymerase, 2 μl barcoding primers). The thermal cycling program was 15 min 72°C, 45 s 95°C, 22 cycles of 15 s 98°C; 30 s 67°C; 1 min 72°C; followed by final elongation at 5 min 72°C; 4°C hold. After PCR, 2 μl of each well was pooled and cleaned-up twice using AMPure XP beads (at 1.3X volume).

Smart-seq3 was performed as described above but with 28 PCR cycles to amplify cDNA. Libraries were sequenced as described above on a NextSeq 550.

#### Allelic RNA dilution series

Liver tissue was isolated from 12-week-old male C57BL6/J and CAST/EiJ mice, dissected into 1-2mm^2^-wide samples and transferred to 1 ml TRIzol isolation reagent (Thermo Scientific). The samples were thoroughly homogenized in TRIzol using a metallic tissue grinder before proceeding to RNA extraction. RNA extraction was performed according to the manufacturer’s instructions. In brief, 0.2 ml chloroform was added to 1ml of TRIzol and the samples were vigorously shaken. The samples were incubated at room temperature and the RNA-containing upper aqueous phase was isolated and precipitated with 0.5 ml isopropyl alcohol. The samples were incubated at room temperature for 10 min and centrifuged at 12000 x g at 4°C for 10 min. The pellet was washed once with 75% ethanol and the RNA pellet was air-dried for 10 min. The RNA pellets were resuspended in 50 μl of RNase-free water and incubated in a heat block at 55-60°C for 15 min before measuring the concentration using a Nanodrop 2000 spectrophotometer. RNA from pure C57BL6/J and CAST/EiJ strains was combined at varying ratios (0, 12.5, 25, 37.5, 50, 62.5, 75, 87.5 and 100% C57) for a total of 200 pg RNA which was subjected to Smart-seq3 with slight modification from above. Briefly, tagmentation was done using 0.1 μl Nextera XT ATM Tn5 [Illumina] for 10 min, index primers were used as 0.2 μM each and a 0.6:1 AMPure XP bead ratio was used for cleanup.

#### Smart-seq3 data analysis

A hybrid mouse genome index was constructed by N-masking the reference genome (GRCm38_68) for CAST/EiJ SNPs from the Mouse Genomes Project (mgp.v5.merged.snps_all.dbSNP142)^5^ using SNPsplit (0.3.2)^6^. Raw Smart-seq3 data was processed using zUMIs (2.8.0)^7^. Briefly, sample barcodes were filtered and data was aligned with STAR (2.7.2a)^8^ [options: --clip3pAdapterSeq CTGTCTCTTATACACATCT] and reads were assigned to both intron and exon features (Mus_musculus.GRCm38.97.chr.gtf) using FeatureCounts^9^. Next, barcodes were collapsed using 1 hamming distance and gene expression was calculated for both reads and UMIs. Finally, allele-level expression was calculated from the zUMIs output as previously described (github.com/sandberg-lab/Smart-seq3/tree/master/allele_level_expression)^2^. 77 cells were excluded due to low read depth (>3 MADs) in the original experiments and 486 cells were excluded due to low read depth in the multi-omics experiments.

#### Processing of published C57BL/6J × CAST/EiJ scRNA-seq data

Pre-processed Smart-seq data for MII oocyte – 16-cell stages were obtained^10^ and two cells were excluded (8cell_8-3 & 16cell_4-2) due to potential sample problems (high paternal X ratio in male and non-ZGA with clustering together with Zygote samples, respectively). Pre-processed blastocyst Smart-seq2 data was obtained^10^ and 18 late blastocysts were excluded due to low read depth (>3 MADs). Pre-processed Smart-seq2 data for postimplantation embryos was obtained^1,11^.

#### Genes escaping X inactivation and X-Y homolog gene lists

A list of known mouse escapee genes (*1810030O07Rik, 2010000l03Rik, 2010308F09Rik, 2610029G23Rik, 5530601H04Rik, 6720401G13Rik, Abcd1, Araf, Atp6ap2, BC022960, Bgn, Car5b, D330035K16Rik, D930009K15Rik, Ddx3x, Eif1ax, Eif2s3x, Fam50a, Flna, Ftsj1, Fundc1, Gdi1, Gemin8, Gpkow, Huwe1, Idh3g, Igbp1*, *Ikbkg*, *Kdm5c*, *Kdm6a*, *Lamp2*, *Maged1*, *Mbtps2*, *Med14*, *Mid1*, *Mmgt1*, *Mpp1*, *Msl3*, *Ndufb11*, *Nkap*, *Ogt*, *Pbdc1, Pdha1, Prdx4, Rbm3, Renbp, Sh3bgrl, Shroom4, Sms, Suv39h1, Syap1, Tbc1d25, Timp1, Trap1a, Uba1, Usp9x, Utp14a, Uxt, Xist, Yipf6)* was compiled from previous work^12–15^ and excluded from analyses where specified.

A list of ancestral X-Y homologs was obtained^16^ and sequence similarity of expressed X-Y homolog transcripts was calculated using the nucleotide BLAST webservice^17^ using default settings (performed 2020-03-24). The ancestral homologs had an average sequence similarity of 83% and were detected in a maximum of 2.5% female cells whereas maximum male-specific detection was 98.3%, indicating that X-Y homologs show correct sexspecific mapping.

#### Expression calculations

Allelic reference ratios were calculated as CountsC57/CountsTotal after exclusion of the top 10% expressed X-linked genes to avoid bias from highly expressed X genes. XCI status was determined as active (Xa/Xa), semi-inactivated (Xa/Xs) or fully inactivated (Xa/Xi) for allelic ratios in the intervals (0.4, 0.6), [0.6, 0.9) and [0.9, ∞) and the inverse, respectively. TPM was calculated for gene *i* in cell *j* as TPM_ij_ = (FPKM_i_, /∑ FPKM_j_) × 10^6^ and allelic TPM was calculated by scaling TPM by reference ratios per gene and cell. Relative UMI counts were calculated relative to total UMIs per sample (percentage of total counts). A gene was considered expressed in a dataset/lineage if the average TPM expression was > 0. To obtain a robust estimate for cell-level expression, 20% trimmed means was calculated per cell.

#### Expression ratios

Chromosome:Autosomal expression ratios were calculated for expressed genes (>1 TPM)^18–20^ as relative to median of autosomes after excluding genes that escape XCI. Additionally, a bootstrapping method^12^ was used to account for different number of genes between chromosomes. For bootstrapped ratios, random autosomal gene sets of the same size as the test set were selected as a background, repeated n = 10^3^ times.

Female: male ratios were calculated after exclusion of X escapees as TPM relative to gene average per embryonic day and lineage, as well as for the active X allele for allelic data.

#### Differential expression of scRNA-seq data

For Smart-seq3 data, global count data was size factor normalized using scater (1.12.2)/scran (1.12.1)^21^ and genes expressed in >10% of cells were kept. Differential expression was calculated along days of differentiation as a continuous variable using likelihood ratio tests as implemented in MAST (1.10.0)^22^. Gene set enrichment for mouse GO biological process gene sets obtained from MGI (http://www.informatics.jax.org/downloads/reports/index.html#go; accessed 2020-06-23) was performed on the differential expression model against a bootstrapped (n = 100) control model as implemented in MAST.

#### Dimensionality reduction, clustering and trajectory inference

For Smart-seq3 data, highly variable genes (HVGs) were identified from size-factor-normalized counts using scater/scran and ordered by biological variance and FDR. Data was visualized for top 1,000 HVGs using diffusion maps^23^.

For *in vivo* blastocyst data, HVGs were obtained as explained above and dimensionality reduction was performed for top 1,000 HVGs using UMAP^24^ and cells were Louvain clustered based on top 1,000 HVG ranks using scran. For *in vivo* post-implantation data, pseudotime trajectories were inferred using Slingshot (1.2.0)^25^ from normalized counts following Mclust (5.4.5)^26^ clustering on diffusion map coordinates.

#### Kinetics inference

Missing allelic data points were set to 0 if the gene was detected on the other allele. Kinetic parameters were calculated per lineage/genotype/XCI status/growth condition (depending on dataset) using txburst (github.com/sandberg-lab/txburst)^27^. Genes not passing filtering steps were excluded and relative burst frequency (k_on_) and burst size (k_syn_/k_off_) was calculated relative to median of expressed autosomal genes (per lineage). Only groups with > 20 cells were kept to increase the reliability of the statistical inference.

#### scATAC data analysis

Raw fastq files were tagged with cell names using GNU sed (4.4), quality and adapter trimmed using fastp (0.20.0)^28^ and aligned to the N-masked reference genome using bowtie2 (2.4.1)^29^ [options: --very-sensitive -N 1 - X 2000 -k 10]. Duplicates were marked using biobambam2 (2.0.87; gitlab.com/german.tischler/biobambam2) and data was merged and split into respective alleles using SNPsplit (0.3.2)^6^ [options: --paired --no_sort]. Downstream processing and analysis was performed using ArchR (1.0.1)^30^. Briefly, non-duplicate properly paired and mapped primary alignments were loaded and mitochondrial and Y chromosomes were excluded. Non-allelic data was filtered for cells with at least 1,000 fragments and a minimum TSS enrichment of 4 and predicted doublets were excluded for a total of 437 excluded cells. Enriched peaks were called grouped by differentiation day using the ArchR implementation of Macs2^31^. Expression data was integrated using Smart-seq3 UMI counts, and dimensionality reduction was performed using the ArchR LSI implementation (default parameters for scATAC and top 1,000 variable features based on variance-to-mean ratio for scRNA) and visualized using UMAP with 15 nearest neighbors. Accessibility X:A ratios was calculated as mean_chrX_/mean_Autosome_ on ArchR gene scores to account for the binary nature of scATAC-seq. Allelic data was filtered to match the cells in the non-allelic data and allelic count matrices were recalculated. Allelic ratios per cell was calculated using the paired expression data, as described above, and X-inactivation completion was calculated per modality as |0.5– (Counts_C57_/Counts_Total_)|/0.5. A list of known imprinted mouse genes (*Sgce, Peg3, Peg12, Plagl1, Zrsr1, Peg13, Airn, Impact, Nckap5, Dlx5, Gm5422, Grb10, Meg3, Rian, Mirg, Igf2r, Igf2, H19*) was obtained^13^ and only genes detected by both scRNA-seq and scATAC-seq was kept for analysis.

#### Re-analysis of published data

Raw bulk RNA-seq data including both male and female mESCs was obtained^32,33^ and quantified to protein-coding transcripts from GENCODE vM22^34^ using pseudoalignment with Salmon (0.14.1)^35^. Transcript abundance estimates were summarized to gene-level using tximport (1.12.3)^36^ and differential expression was calculated using likelihood ratio tests in DESeq2 (1.24.0)^37^. A full model including cellular genotype (XX, XY or XO), cell culture condition (2i or serum) and study accession was tested against a reduced model without the genotype term. Lowly expressed genes (<100 average normalized counts) were excluded from final plots. Serum growth conditions only showed a minor effect on global X expression compared to 2i (not shown), consistent with a low degree of cells exhibiting complete XCI^1^.

Pre-processed scRNA-seq data for wild-type and *Xist^patΔ^* knockout embryos was obtained^12^; processed data from GSE80810. Briefly, allelic reference ratios were used to split normalized expression values (RPRT; Reads Per Retro-Transcribed length per million mapped reads) into alleles and average expression per cell was calculated using trimmed means for expressed genes (average total RPRT > 0) as described above.

Pre-processed scRNA-seq data for mESCs adapting to serum/LIF conditions were obtained^38^; processed data from GSE151009. Briefly, Total UMI count matrices were size factor normalized as described above and allelic UMI count matrices were used as is. Reference ratios and XCI status was calculated as described above.

Raw native ChIP-seq data for hybrid mESCs was obtained^39^, quality and adapter trimmed using fastp (0.20.0)^28^, aligned to a CAST/EiJ N-masked genome using Bowtie2 (2.4.1)^29^ [options: -N 1] and aligned data was split to respective alleles using SNPsplit (0.3.2)^6^. Peaks were called against Input samples as controls using MACS2 (2.1.2)^31^ [options: --broad -broad-cutof 0.01 -f BAMPE -g 2652783500], peaks overlapping ENCODE v2 problematic regions^40^ were excluded and consensus peaks were defined as peaks with a reciprocal overlap of at least 25% of peak width between replicates. Normalized coverage was calculated using deepTools (3.3.0)^41^[options bamCoverage –normalizeUsing RPGC –effectiveGenomeSize 2652783500 --skipNAs --ignoreDuplicates --centerReads --blackListFileName mm10-blacklist.v2.ENSEMBL.bed.gz] and profiles were calculated using deepTools for chrX TSSs (Mus_musculus.GRCm38.97.chr.gtf) [options: computeMatrix reference-point --upstream 5000 --downstream 5000 --skipZeros --nanAfterEnd] or chrX enhancers (intersection between H3K4me1 and H3K27ac consensus peaks excluding peaks withing 1kb of a TSS) [options: computeMatrix reference-point --referencePoint center --upstream 5000 --downstream 5000 --skipZeros] and resulting coverage profiles were normalized against Input.

Raw *in situ* Hi-C data for hybrid mESCs was obtained^42^. An N-masked dual hybrid mouse genome index was constructed for 129S1/svImJ x CAST/EiJ (mgp.v5.merged.snps_all.dbSNP142) using SNPsplit (0.3.2)^6^ and further *in silico* MboI digested for downstream tools. Data was aligned using HiCUP (0.8.0) [options: --bowtie2 --shortest 50 --longest 700 --digest –zip]^43^ then split into alleles using SNPsplit, converted to juicer format using HiCUP (hicup2juicer) and reads uniquely corresponding to either allele (requiring at least one mate to map to the genome) were merged. Hi-C contact maps were generated using juicer tools pre (1.22.01) [options: -d -f -q 10 -r 1000000]^44^ and KR-normalized 1Mb matrices were extracted using juicer tools (dump observed/pearsons/eigenvector). Observed contact counts were further normalized for sequencing depth (per million) per chromosome and timepoint.

#### Statistics and data visualization

All statistical tests were performed in R (3.6.1) as two-tailed unless otherwise stated. Heatmaps were visualized using ComplexHeatmap (2.0.0)^45^ and all other plots were made using ggplot2 (3.2.1)^46^. Box plots are presented as median, first and third quartiles, and 1.5x inter-quartile range (IQR). For median ± confidence interval plots, bootstrapped 95% confidence intervals (n = 1,000) were calculated using the percentile method^47^ as implemented in the R boot package (1.3-23).

**Other Supplementary Materials for this manuscript includes the following:**

**Supplementary Table 1 (Microsoft Excel format).** Differentially expressed genes along mESC differentiation towards EpiSCs and gene ontology (GO) term enrichment.

**Supplementary Table 2 (Microsoft Excel format).** Sequence identity of X-Y gene pairs.

## References

1. Deng, X., Berletch, J. B., Nguyen, D. K. & Disteche, C. M. X chromosome regulation: Diverse patterns in development, tissues and disease. Nat. Rev. Genet. 15, 367–378 (2014).

2. Ohno, S. Sex Chromosomes and Sex-linked Genes. In Monographs on endocrinology. Springer-Verl. Heidelb.-Berl.-N. Y. 1, (1967).

3. Graves, J. A. M. Sex chromosome specialization and degeneration in mammals. Cell 124, 901–914 (2006).

4. Gribnau, J. & Grootegoed, J. A. Origin and evolution of X chromosome inactivation. Curr. Opin. Cell Biol. 24, 397–404 (2012).

5. Okamoto, I., Otte, A. P., Allis, C. D., Reinberg, D. & Heard, E. Epigenetic dynamics of imprinted X inactivation during early mouse development. Science 303, 644–649 (2004).

6. Deng, Q., Ramsköld, D., Reinius, B. & Sandberg, R. Singlecell RNA-seq reveals dynamic, random monoallelic gene expression in mammalian cells. Science 343, 193–196 (2014).

7. Deuve, J. L. & Avner, P. The Coupling of X-Chromosome Inactivation to Pluripotency. Annu. Rev. Cell Dev. Biol. 27, 611–629 (2011).

8. Nguyen, D. K. & Disteche, C. M. Dosage compensation of the active X chromosome in mammals. Nat. Genet. 38, 47–53 (2006).

9. Gupta, V. et al. Global analysis of X-chromosome dosage compensation. J. Biol. 5, (2006).

10. Lin, H. et al. Dosage compensation in the mouse balances upregulation and silencing of X-linked genes. PLoS Biol. 5, 2809–2820 (2007).

11. Deng, X. et al. Evidence for compensatory upregulation of expressed X-linked genes in mammals, Caenorhabditis elegans and Drosophila melanogaster. Nat. Genet. 43, 1179–1185 (2011).

12. Deng, X. et al. Mammalian X upregulation is associated with enhanced transcription initiation, RNA half-life, and MOF-mediated H4K16 acetylation. Dev. Cell 25, 55–68 (2013).

13. Mahadevaiah, S. K., Sangrithi, M. N., Hirota, T. & Turner, J. M. A. A single-cell transcriptome atlas of marsupial embryogenesis and X inactivation. Nature (2020) doi:10.1038/s41586-020-2629-6.

14. Borensztein, M. et al. Xist-dependent imprinted X inactivation and the early developmental consequences of its failure. Nat. Struct. Mol. Biol. 24, 226–233 (2017).

15. Wang, F. et al. Regulation of X-linked gene expression during early mouse development by Rlim. eLife 5, (2016).

16. Reinius, B. & Sandberg, R. Random monoallelic expression of autosomal genes: stochastic transcription and allele-level regulation. Nat. Rev. Genet. 16, 653–664 (2015).

17. Chen, G. et al. Single-cell analyses of X Chromosome inactivation dynamics and pluripotency during differentiation. Genome Res. 26, 1342–1354 (2016).

18. Hagemann-Jensen, M. et al. Single-cell RNA counting at allele and isoform resolution using Smart-seq3. Nat. Biotechnol. 38, 708–714 (2020).

19. Marks, H. et al. Dynamics of gene silencing during X inactivation using allele-specific RNA-seq. Genome Biol. 16, 9–10 (2015).

20. Werner, R. J. et al. Sex chromosomes drive gene expression and regulatory dimorphisms in mouse embryonic stem cells. Biol. Sex Differ. 8, 28 (2017).

21. Pacini, G. et al. Integrated analysis of Xist upregulation and X-chromosome inactivation with single-cell and single-allele resolution. Nat. Commun. 12, 3638 (2021).

22. Schulz, E. G. et al. The Two Active X Chromosomes in Female ESCs Block Exit from the Pluripotent State by Modulating the ESC Signaling Network. Cell Stem Cell 14, 203–216 (2014).

23. Cheng, S. et al. Single-Cell RNA-Seq Reveals Cellular Heterogeneity of Pluripotency Transition and X Chromosome Dynamics during Early Mouse Development. Cell Rep. 26, 2593–2607.e3 (2019).

24. Deng, X. & Disteche, C. M. Rapid transcriptional bursts upregulate the X chromosome. Nat. Struct. Mol. Biol. 26, 851–853 (2019).

25. Larsson, A. J. M., Coucoravas, C., Sandberg, R. & Reinius, B. X-chromosome upregulation is driven by increased burst frequency. Nat. Struct. Mol. Biol. 26, 963–969 (2019).

26. Carrel, L. & Willard, H. F. X-inactivation profile reveals extensive variability in X-linked gene expression in females. Nature 434, 400–404 (2005).

27. Yang, F., Babak, T., Shendure, J. & Disteche, C. M. Global survey of escape from X inactivation by RNA-sequencing in mouse. Genome Res. 20, 614–622 (2010).

28. Tukiainen, T. et al. Landscape of X chromosome inactivation across human tissues. Nature 550, 244–248 (2017).

29. Soh, Y. Q. S. et al. Sequencing the mouse y chromosome reveals convergent gene acquisition and amplification on both sex chromosomes. Cell 159, 800–813 (2014).

30. Sangrithi, M. N. et al. Non-Canonical and Sexually Dimorphic X Dosage Compensation States in the Mouse and Human Germline. Dev. Cell 40, 289–301.e3 (2017).

31. Yildirim, E., Sadreyev, R. I., Pinter, S. F. & Lee, J. T. X-chromosome hyperactivation in mammals via nonlinear relationships between chromatin states and transcription. Nat. Struct. Mol. Biol. 19, 56–62 (2012).

32. Larsson, A. J. M. et al. Genomic encoding of transcriptional burst kinetics. Nature 565, 251–254 (2019).

33. Reinius, B. et al. Analysis of allelic expression patterns in clonal somatic cells by single-cell RNA-seq. Nat. Genet. 48, 1430–1435 (2016).

34. Zylicz, J. J. et al. The Implication of Early Chromatin Changes in X Chromosome Inactivation. Cell 176, 182–197.e23 (2019).

35. Bartman, C. R., Hsu, S. C., Hsiung, C. C.-S., Raj, A. & Blobel, G. A. Enhancer Regulation of Transcriptional Bursting Parameters Revealed by Forced Chromatin Looping. Mol. Cell 62, 237–247 (2016).

36. Fukaya, T., Lim, B. & Levine, M. Enhancer Control of Transcriptional Bursting. Cell 166, 358–368 (2016).

37. Froberg, J. E., Pinter, S. F., Kriz, A. J., Jégu, T. & Lee, J. T. Megadomains and superloops form dynamically but are dispensable for X-chromosome inactivation and gene escape. Nat. Commun. 9, 5004 (2018).

38. Deng, X. et al. Bipartite structure of the inactive mouse X chromosome. Genome Biol. 16, 152 (2015).

39. Giorgetti, L. et al. Structural organization of the inactive X chromosome in the mouse. Nature 535, 575–579 (2016).

40. Bauer, M. et al. Chromosome compartments on the inactive X guide TAD formation independently of transcription during X-reactivation. Nat. Commun. 12, 3499 (2021).

41. Wang, M., Lin, F., Xing, K. & Liu, L. Random X-chromosome inactivation dynamics in vivo by single-cell RNA sequencing. BMC Genomics 18, 9–10 (2017).

42. Fukuda, A., Tanino, M., Matoba, R., Umezawa, A. & Akutsu, H. Imbalance between the expression dosages of X-chromosome and autosomal genes in mammalian oocytes. Sci. Rep. 5, 9–10 (2015).

43. Li, X. et al. Dosage compensation in the process of inactivation/reactivation during both germ cell development and early embryogenesis in mouse. Sci. Rep. 7, (2017).

44. Chitiashvili, T. et al. Female human primordial germ cells display X-chromosome dosage compensation despite the absence of X-inactivation. Nat. Cell Biol. 22, 1436–1446 (2020).

45. Takagi, N. & Abe, K. Detrimental effects of two active X chromosomes on early mouse development. Dev. Camb. Engl. 109, 189–201 (1990).

46. Cortini, R. & Filion, G. J. Theoretical principles of transcription factor traffic on folded chromatin. Nat. Commun. 9, 9–10 (2018).

47. Brouwer, I. & Lenstra, T. L. Visualizing transcription: key to understanding gene expression dynamics. Curr. Opin. Chem. Biol. 51, 122–129 (2019).

48. Chaumeil, J. A novel role for Xist RNA in the formation of a repressive nuclear compartment into which genes are recruited when silenced. Genes Dev. 20, 2223–2237 (2006).

49. Fan, G. et al. X-chromosome dosage compensation dynamics in human early embryos. bioRxiv 2020.03.08.982694 (2020) doi:10.1101/2020.03.08.982694.

50. Augui, S., Nora, E. P. & Heard, E. Regulation of X-chromosome inactivation by the X-inactivation centre. Nat. Rev. Genet. 12, 429–442 (2011).

## Supplementary references

1. Chen, G. et al. Single-cell analyses of X Chromosome inactivation dynamics and pluripotency during differentiation. Genome Research 26, 1342–1354 (2016).

2. Hagemann-Jensen, M. et al. Single-cell RNA counting at allele and isoform resolution using Smart-seq3. Nat Biotechnol 38, 708–714 (2020).

3. Picelli, S. et al. Tn5 transposase and tagmentation procedures for massively scaled sequencing projects. Genome Res. 24, 2033–2040 (2014).

4. Zachariadis, V., Cheng, H., Andrews, N. & Enge, M. A Highly Scalable Method for Joint Whole-Genome Sequencing and Gene-Expression Profiling of Single Cells. Molecular Cell 80, 541–553.e5 (2020).

5. Keane, T. M. et al. Mouse genomic variation and its effect on phenotypes and gene regulation. Nature 477, 289–294 (2011).

6. Krueger, F. & Andrews, S. R. SNPsplit: Allele-specific splitting of alignments between genomes with known SNP genotypes. F1000Res 5, 1479 (2016).

7. Parekh, S., Ziegenhain, C., Vieth, B., Enard, W. & Hellmann, I. zUMIs - A fast and flexible pipeline to process RNA sequencing data with UMIs. GigaScience 7, (2018).

8. Dobin, A. et al. STAR: ultrafast universal RNA-seq aligner. Bioinformatics 29, 15–21 (2013).

9. Liao, Y., Smyth, G. K. & Shi, W. The R package Rsubread is easier, faster, cheaper and better for alignment and quantification of RNA sequencing reads. Nucleic Acids Research 47, e47–e47 (2019).

10. Deng, Q., Ramsköld, D., Reinius, B. & Sandberg, R. Single-cell RNA-seq reveals dynamic, random monoallelic gene expression in mammalian cells. Science 343, 193–196 (2014).

11. Cheng, S. et al. Single-Cell RNA-Seq Reveals Cellular Heterogeneity of Pluripotency Transition and X Chromosome Dynamics during Early Mouse Development. Cell Reports 26, 2593–2607.e3 (2019).

12. Borensztein, M. et al. Xist-dependent imprinted X inactivation and the early developmental consequences of its failure. Nature Structural and Molecular Biology 24, 226–233 (2017).

13. Reinius, B. et al. Analysis of allelic expression patterns in clonal somatic cells by single-cell RNA-seq. Nat Genet 48, 1430–1435 (2016).

14. Reinius, B. et al. Female-biased expression of long non-coding RNAs in domains that escape X-inactivation in mouse. BMC Genomics 11, (2010).

15. Yang, F., Babak, T., Shendure, J. & Disteche, C. M. Global survey of escape from X inactivation by RNA-sequencing in mouse. Genome Research 20, 614–622 (2010).

16. Soh, Y. Q. S. et al. Sequencing the mouse y chromosome reveals convergent gene acquisition and amplification on both sex chromosomes. Cell 159, 800–813 (2014).

17. Johnson, M. et al. NCBI BLAST: a better web interface. Nucleic Acids Res. 36, W5–9 (2008).

18. Deng, X. et al. Evidence for compensatory upregulation of expressed X-linked genes in mammals, Caenorhabditis elegans and Drosophila melanogaster. Nature Genetics 43, 1179–1185 (2011).

19. Lin, H. et al. Dosage compensation in the mouse balances up-regulation and silencing of X-linked genes. PLoS Biology 5, 2809–2820 (2007).

20. Yildirim, E., Sadreyev, R. I., Pinter, S. F. & Lee, J. T. X-chromosome hyperactivation in mammals via nonlinear relationships between chromatin states and transcription. Nature Structural and Molecular Biology 19, 56–62 (2012).

21. Lun, A. T. L., McCarthy, D. J. & Marioni, J. C. A step-by-step workflow for low-level analysis of single-cell RNA-seq data with Bioconductor. F1000Research 5, 2122 (2016).

22. Finak, G. et al. MAST: A flexible statistical framework for assessing transcriptional changes and characterizing heterogeneity in single-cell RNA sequencing data. Genome Biology 16, 9–10 (2015).

23. Haghverdi, L., Buettner, F. & Theis, F. J. Diffusion maps for high-dimensional single-cell analysis of differentiation data. Bioinformatics 31, 2989–2998 (2015).

24. McInnes, L., Healy, J. & Melville, J. UMAP: Uniform Manifold Approximation and Projection for Dimension Reduction. (2018).

25. Street, K. et al. Slingshot: Cell lineage and pseudotime inference for single-cell transcriptomics. BMC Genomics 19, (2018).

26. Fraley, C. & Raftery, A. E. Bayesian regularization for normal mixture estimation and model-based clustering. Journal of Classification 24, 155–181 (2007).

27. Larsson, A. J. M. et al. Genomic encoding of transcriptional burst kinetics. Nature 565, 251–254 (2019).

28. Chen, S., Zhou, Y., Chen, Y. & Gu, J. fastp: an ultra-fast all-in-one FASTQ preprocessor. Bioinformatics 34, i884–i890 (2018).

29. Langmead, B. & Salzberg, S. L. Fast gapped-read alignment with Bowtie 2. Nat Methods 9, 357–359 (2012).

30. Granja, J. M. et al. ArchR is a scalable software package for integrative single-cell chromatin accessibility analysis. Nat Genet 53, 403–411 (2021).

31. Zhang, Y. et al. Model-based Analysis of ChIP-Seq (MACS). Genome Biol 9, R137 (2008).

32. Marks, H. et al. The transcriptional and epigenomic foundations of ground state pluripotency. Cell 149, 590–604 (2012).

33. Werner, R. J. et al. Sex chromosomes drive gene expression and regulatory dimorphisms in mouse embryonic stem cells. Biol Sex Differ 8, 28 (2017).

34. Frankish, A. et al. GENCODE reference annotation for the human and mouse genomes. Nucleic Acids Research 47, D766–D773 (2019).

35. Patro, R., Duggal, G., Love, M. I., Irizarry, R. A. & Kingsford, C. Salmon provides fast and bias-aware quantification of transcript expression. Nat Methods 14, 417–419 (2017).

36. Soneson, C., Love, M. I. & Robinson, M. D. Differential analyses for RNA-seq: transcript-level estimates improve genelevel inferences. F1000Res 4, 1521 (2015).

37. Love, M. I., Huber, W. & Anders, S. Moderated estimation of fold change and dispersion for RNA-seq data with DESeq2. Genome Biol 15, 550 (2014).

38. Pacini, G. et al. Integrated analysis of Xist upregulation and X-chromosome inactivation with single-cell and single-allele resolution. Nat Commun 12, 3638 (2021).

39. Zylicz, J. J. et al. The Implication of Early Chromatin Changes in X Chromosome Inactivation. Cell 176, 182–197.e23 (2019).

40. Amemiya, H. M., Kundaje, A. & Boyle, A. P. The ENCODE Blacklist: Identification of Problematic Regions of the Genome. Sci Rep 9, 9354 (2019).

41. Ramírez, F. et al. deepTools2: a next generation web server for deep-sequencing data analysis. Nucleic Acids Res 44, W160–W165 (2016).

42. Froberg, J. E., Pinter, S. F., Kriz, A. J., Jégu, T. & Lee, J. T. Megadomains and superloops form dynamically but are dispensable for X-chromosome inactivation and gene escape. Nat Commun 9, 5004 (2018).

43. Wingett, S. W. et al. HiCUP: pipeline for mapping and processing Hi-C data. F1000Res 4, 1310 (2015).

44. Durand, N. C. et al. Juicer Provides a One-Click System for Analyzing Loop-Resolution Hi-C Experiments. Cell Systems 3, 95–98 (2016).

45. Gu, Z., Eils, R. & Schlesner, M. Complex heatmaps reveal patterns and correlations in multidimensional genomic data. Bioinformatics 32, 2847–2849 (2016).

46. Wickham, H. Ggplot2: Elegant Graphics for Data Analysis. vol. 174 (2016).

47. Davison, A. C. & Hinkley, D. V. Bootstrap Methods and their Application. Bootstrap Methods and their Application (1997) doi:10.1017/cbo9780511802843.

